# Restricting α-Synuclein Transport into Mitochondria by Inhibition of α-Synuclein-VDAC Complexation as a Potential Therapeutic Target for Parkinson’s Disease Treatment

**DOI:** 10.1101/2022.04.07.487541

**Authors:** Megha Rajendran, María Queralt-Martín, Philip A. Gurnev, William M. Rosencrans, Amandine Rovini, Daniel Jacobs, Kaitlin Abrantes, David P. Hoogerheide, Sergey M. Bezrukov, Tatiana K. Rostovtseva

## Abstract

Involvement of alpha-synuclein (αSyn) in Parkinson’s disease (PD) is complicated and difficult to trace on cellular and molecular levels. Recently we established that αSyn can regulate mitochondrial function by voltage-activated complexation with the Voltage-Dependent Anion Channel (VDAC) of the outer mitochondrial membrane. When complexed with αSyn, the VDAC pore is partially blocked, reducing the transport of ATP/ADP and other metabolites. Further, αSyn can translocate into the mitochondria through VDAC, where it interferes with mitochondrial respiration. Recruitment of αSyn to the VDAC-containing lipid membrane appears to be a crucial prerequisite for both the blockage and translocation processes. Here we report an inhibitory effect of HK2p, a small membrane-binding peptide from the mitochondria-targeting N-terminus of hexokinase 2, on the αSyn membrane binding, and hence on αSyn complex formation with VDAC and translocation through it. In electrophysiology experiments, addition of HK2p at micromolar concentrations to the same side of the membrane as αSyn results in dramatic reduction of the frequency of blockage events in a concentration-dependent manner, reporting on complexation inhibition. Using two complementary methods of measuring protein-membrane binding, bilayer overtone analysis and fluorescence correlation spectroscopy, we found that HK2p induces detachment of αSyn from lipid membranes. Experiments with live HeLa cells using proximity ligation assay confirmed that HK2p impedes αSyn entry into mitochondria. Our results demonstrate that it is possible to regulate αSyn-VDAC complexation by a rationally designed peptide, thus suggesting new avenues in the search for peptide therapeutics to alleviate αSyn mitochondrial toxicity in PD and other synucleinopathies.

## INTRODUCTION

The intrinsically disordered neuronal protein α-synuclein (αSyn) is found accumulated in a fibrillar form in Lewy bodies [1], a hallmark of Parkinson’s disease (PD) [2]. Nevertheless, understanding the role of αSyn in PD remains challenging. The general paradigm centers on the aggregation of overproduced αSyn into progressively larger amyloids that cause cell toxicity [3], with other aspects of αSyn cellular distribution and function remaining unclear. The role of αSyn in mitochondria and certain features of underlying mitochondrial dysfunction in PD have been suggested previously [3–8]. Overproduction of αSyn is known to cause serious consequences such as loss of mitochondrial potential, decrease of ATP level, increase of reactive oxygen species (ROS) production, and impairment of the electron transport chain (ETC) complexes [7–9]. However, to target ETC complexes at the mitochondrial inner membrane (IM), the water-soluble cytosolic αSyn must first cross the mitochondrial outer membrane (MOM) through a specific membrane transport system such as voltage-dependent anion channel (VDAC). One of the likely causes of αSyn pathological effects is the mitochondrial toxicity of its monomeric, rather than oligomeric, form through its interaction with VDAC[10–12].

Using single-channel electrophysiology experiments, it was recently shown that at nanomolar solution concentrations, monomeric αSyn reversibly blocks VDAC reconstituted into planar lipid membranes in a concentration- and voltage-dependent manner and, importantly, can translocate through this channel [10, 13]. Therefore, it was suggested that αSyn could regulate the transport of metabolites (ATP, ADP, etc.) and Ca^2+^ through the VDAC pore and that αSyn translocates through this pore into mitochondria [11, 12]. Under certain conditions, such as αSyn overexpression, increased VDAC blockage and translocation of αSyn into mitochondria may result in mitochondrial dysfunction seen in PD. Experiments in cells supported the electrophysiological results by first demonstrating that VDAC is required for αSyn toxicity using the yeast strain deficient in VDAC1, in which human VDAC1 and wild type (WT) αSyn were overexpressed [10]. Later, using an *in situ* Proximity Ligation Assay (PLA) on a relevant PD cell model of human differentiated dopaminergic neuronal cell line with overexpressed WT αSyn, αSyn was found in close proximity to VDAC in the MOM and complex IV in the IM [11]. Knockdown of VDAC1 using siRNA resulted in the reduction of αSyn transport into the mitochondria, thus adding further evidence that VDAC forms a pathway for αSyn translocation across the MOM and is needed for αSyn-induced mitochondrial toxicity [11]. As VDAC controls fluxes of metabolites in and out of mitochondria, these findings may represent the mechanisms of both the regulatory role of monomeric αSyn and its toxicity to mitochondria.

αSyn is a relatively small, 140-residue polypeptide that is disordered in solution but adopts an α-helical structure when bound to anionic lipid membranes [14–16]. αSyn consists of three distinct domains: an N-terminal amphipathic membrane-binding domain, a central nonpolar or so-called non-amyloid-ß component (NAC) domain, which is involved in fibril formation, and a highly negatively charged disordered C-terminal tail (CTT) domain [15]. A combination of electrophysiological and biophysical methods along with theoretical modeling demonstrates that αSyn-VDAC complexation occurs by a two-step mechanism [10, 17–19]. First, αSyn is recruited to a lipid membrane, binding to the lipids by its amphipathic N-terminal domain. Second, complex formation proceeds by the voltage-induced capture of the negatively charged CTT of αSyn by the VDAC pore. The exclusion of small, current-carrying ions from the pore when the CTT is inserted leads to transient ionic current blockages [19] (Figure 1A, *steps 1-3* and Figure 1B, *two left traces*). Eventually, the αSyn-VDAC complex dissociates in one of two ways: at small applied potentials, the membrane-bound N-terminal domain of αSyn prevents translocation of the entire molecule through the channel, leading to eventual withdrawal, or retraction, of the CTT from the VDAC pore back to the same side of the channel where αSyn entered it. At relatively high potentials, on the other hand, the CTT can be pulled hard enough to release the N-terminal domain from the membrane, leading to αSyn translocation through the pore [10, 17]. This mechanism of the voltage-driven αSyn-VDAC complexation highlights the membrane recruitment of αSyn as a key step in synuclein translocation into mitochondria [17, 19, 20] and, therefore, points to a promising target for specifically designed drugs.

**Figure 1.**
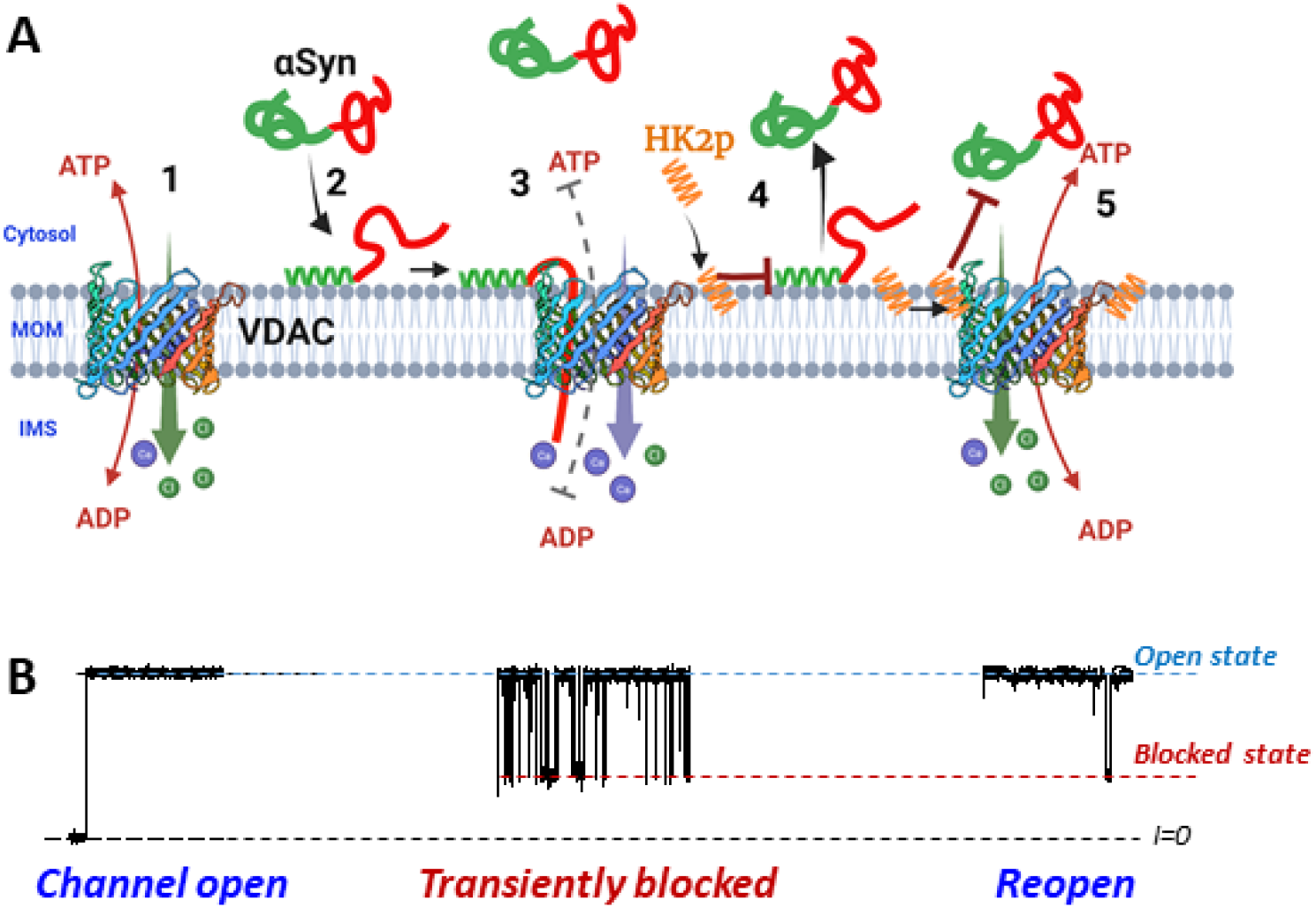
A proposed model of HK2p effect on VDAC permeability. (**A**) The cartoon illustrates how a competitor to αSyn binding to the MOM could counteract αSyn-VDAC complexation and promote metabolite fluxes through the channel. Open VDAC (step 1) maintains effective ATP/ADP transport and low Ca^2+^ permeability. αSyn binds to the membrane by its N-terminal domain (shown in green) (step 2). This binding is a prerequisite to αSyn capture by the pore (step 3). Anionic CTT of αSyn (shown in red) detained in the pore reverses VDAC selectivity to cationic causing reduction of ATP/ADP transport and increase of Ca^2+^ permeability. Hydrophobic HK2p competes with αSyn at the membrane (step 4) inhibiting VDAC blockage and αSyn translocation (step 5). Created with BioRender.com. (**B**) Corresponding VDAC current traces. The exclusion of small, current-carrying ions from the pore when the CTT of αSyn is inserted leads to transient ionic current blockages (middle trace)

Here, we disrupt the membrane recruitment of αSyn using a small synthetic peptide which competes with αSyn for lipid membrane binding sites and is thus expected to downregulate the VDAC-facilitated transport of αSyn (Figure 1A, *steps 4-5*). For this purpose, we choose HK2p, a peptide comprising the mitochondria-targeting domain (residues 1-15) of the hexokinase 2 protein, based on its well-known ability to selectively target mitochondria [21–24] and its reported direct interaction with VDAC [25–27]. We show that HK2p, added in micromolar concentrations to the same side of the VDAC-containing planar membrane as αSyn, reduces the frequency of αSyn-induced VDAC blockage events in a dose-dependent manner. By using two complementary biophysical methods, Bilayer Overtone Analysis (BOA) and Fluorescence Correlation Spectroscopy (FCS), we find that HK2p induces αSyn desorption from the membrane surface, independent of VDAC’s presence in the membrane. We thus validate that competition for the lipid binding sites with αSyn is the underlying cause of HK2p’s effect on αSyn-VDAC complexation. Using a cell-penetrating HK2p construct and *in situ* PLA on HeLa cells with endogenous αSyn, we show that HK2p reduces αSyn entry into mitochondria. These results demonstrate that it is possible to regulate αSyn complexation with VDAC and translocation through it by a rationally chosen small peptide, thus opening new avenues in the search for small-peptide inhibitors of αSyn mitochondrial toxicity in PD and other synucleinopathies. Targeting αSyn association with the outer surface of mitochondrial membrane and αSyn transport into these energy-producing organelles could be a viable but previously unexplored opportunity to reduce αSyn toxicity and to protect mitochondria.

## MATERIALS AND METHODS

### Peptide synthesis

The mitochondria membrane binding peptide sequence from human hexokinase-2, MIASHLLAYFFTELN, was linked to cell-penetrating peptides (CPP): HK2p-CPP(R9) and HK2p-CPP (TAT) for cell penetration and synthesized by Genscript, Inc Control peptide ScrP-CPP was synthesized by Peptide2.0. Labeled peptides with N-terminal tetramethyl Rhodamine (TMR): TMR-HK2p-CPP and control TMR-ScrP-CPP were synthesized by GenScript, Inc. C-terminal TMR labeled, HK2p-CPP-TMR and ScrP-CPP-TMR were synthesized by Alan Scientific, Inc and Genscript, Inc, respectively.

**Table 1.** Synthetic constructs.

| Peptide | Sequence |
| --- | --- |
| HK2p | MIASHLLAYFFTELN |
| HK2p-CPP (R9) | MIASHLLAYFFTELNRRRRRRRRR |
| ScrP-CPP (R9) | VGAHAGEYGAEALER-βA-RRRRRRRRR |
| HK2p-CPP (TAT) | MIASHLLAYFFTELN-βA-GYGRKKRRQRRRG |
| TMR-HK2p-CPP | TMR-MIASHLLAYFFTELN-βA-RRRRRRRRR |
| TMR-ScrP-CPP | TMR-VGAHAGEYGAEALER-βA-RRRRRRRRR |
| HK2p-CPP-TMR | MIASHLLAYFFTELN-βA-RRRRRRRRR-Lys(TMR) |
| ScrP-CPP- TMR | VGAHAGEYGAEALER-βA-RRRRRRRRR-Lys(TMR) |

HK2p was dissolved in N-Methyl-2-pyrrolidone (MP) (Sigma) for channel experiments. Covalently linking CPP to HK2p and ScrP increases their water solubility.

### Protein Purification

VDAC was isolated from frozen mitochondrial fractions of rat liver that were a kind gift of Dr. Marco Colombini (University of Maryland, College Park, MD) following the standard method (Blachly-Dyson et al Science, 1990), and purified on a hydroxyapatite/celite (2:1) column as described previously (Palmieri, De Pinto,1989). VDAC purified from the mitochondrial fraction of the liver contains all three isoforms with VDAC1 being the predominant one (~ 80% of total VDACs) (Yamamoto et al, J Proteome Res 2006). Recombinant WT and Alexa-488-labeled αSyn were the generous gifts of Dr. Jennifer Lee (NHLBI, NIH). WT αSyn and its Cys variant (Y136C) were expressed, purified, and characterized as described previously [28] and stored at −80 °C. αSyn Y136C mutant was labeled with Alexa Fluor 488 at position Y136C as described previously [20]. WT α-hemolysin (Calbiochem, Merck, Darmstadt, Germany) was diluted in an aqueous solution containing 8 M urea at 25 μg/mL (M = mol/L).

### Lipids

1,2-dioleoyl-sn-glycero-3-phosphocholine (DOPC), 1,2-dioleoyl-sn-glycero-3-phosphoethanolamine (DOPE), 1,2-dioleoyl-sn-glycero-3-phospho-(1’-rac-glycerol) (DOPG), diphytanoyl phosphatidylcholine (DPhPC), a soybean Polar Lipid Extract (PLE), and cholesterol (chol) were purchased from Avanti Polar Lipids (Alabaster, AL). All other chemicals were obtained from Sigma-Aldrich unless noted otherwise.

### Live cell imaging

For imaging HK2p-CPP targeting to the mitochondria, HeLa cells were grown in 8-well chamber coverslip (80826, Ibidi USA, Inc) for 24 hours in DMEM (10567, Gibco) supplemented with 10% FBS (F0926, Sigma Aldrich). HeLa cells were either transiently transfected with mito-GFP using LipoSTEM (STEM0008, Thermofisher) the day before imaging or incubated with mitotracker green FM (M7514, Thermofisher) for 15 mins at 37 °C to label the mitochondria. The cells were washed with DPBS (14040, Gibco) and incubated with 5 μM of TMR-labeled peptides for 10 mins at room temperature (RT). The cells were washed and imaged in Fluorobrite DMEM (A18967, Gibco) using a Leica TCS SP8 microscope with a plan-apochromat 100x/ 1.40 N.A. oil objective. Mitotracker green or mito-GFP was visualized by exciting with 488 nm lasers and emission captured using 490-535nm filter and TMR labeled peptides were excited with 552 nm lasers and emission captured using 558-715nm filter. Images were optimized for contrast and brightness using ImageJ (NIH).

### Proximity ligation assay (PLA)

HeLa cells were grown on 12 well-chamber slides (81201, Ibidi USA, Inc). PLA was carried out using Duolink In Situ reagent red (DUO92101, Sigma Aldrich) with a few modifications described here. The cells were treated with 5 μM HK2p-CPP (R9) or Scrp-CPP (R9) for 2-10 min at room temperature (RT). The cells were then fixed with 3.7% paraformaldehyde (15710, Electron microscopy) for 10 min. The cells were incubated with 50 mM glycine for 20 mins and washed three times with PBS. The cells were permeabilized with 0.1% Triton-X100 in 3% BSA-PBS for 15 mins at RT. After washing with PBS, the cells were incubated with Duolink blocking solution for 60 mins at 37 °C. The cells were then incubated overnight with primary antibodies for αSyn (2628s, Cell Signaling Technology, CST) and CoxIV (11967S, CST) at 4°C. Cells were washed twice with wash buffer A and incubated with anti-rabbit PLA probe plus and anti-mouse PLA probe minus along with Tom20-Alexafluor488 antibody (CL488-66777, Proteintech) to label mitochondria for 1 hour at 37 °C. Cells were then incubated with Duolink ligase solution for 30 mins, followed by Duolink amplification polymerase solution for 100 mins at 37 °C. The cells were finally washed with wash buffer B and mounted with Duolink in situ mounting medium with DAPI (DUO82040, Sigma Aldrich). Cells were imaged with Leica TCS SP8 microscope using plan apochromat 40X/1.3 N.A oil objective. The cells were excited at 405 nm (DAPI), 488 nm (Tom20-Alexa488), and 594 nm (PLA dots), and images were collected using 415-460nm, 497-550 nm, and 600-750 nm filters, respectively. The image analysis was performed as previously described [29].

### Channel reconstitution and conductance measurements

The procedure of VDAC reconstitution into lipid bilayers was described previously [10, 30]. Bilayers were formed from DOPG/DOPC/DOPE (2:1:1) (mol/mol), DPhPC, or PLE/chol/DPhPC (90:5:5) (w/w) mixture in 1 M KCl aqueous solutions buffered with 5 mM HEPES at pH 7.4. Potential is defined as positive when it is greater at the side of VDAC addition (*cis*-side). αSyn at a final concentration of 50 nM was added symmetrically to both sides of the membrane under constant stirring for 2 min after VDAC channel reconstitution; statistical analysis of the blockage events was started 15 min after αSyn addition to ensure a steady state. Conductance measurements were performed as described previously [10] using an Axopatch 200B amplifier (Axon Instruments, Inc., Foster City, CA) in the voltage clamp mode. Data were filtered by a low pass 8-pole Butterworth filter (Model 900, Frequency Devices, Inc., Haverhill, MA) at 15 kHz and a low pass Bessel filter at 10 kHz and directly saved into computer memory with a sampling frequency of 50 kHz. For data analysis by Clampfit 10.3, a digital 8-pole Bessel low pass filter set at 5 kHz was applied to current recordings. Individual events of current blockages were discriminated by a threshold crossing algorithm, while kinetic parameters were acquired by fitting single exponentials to logarithmically binned histograms [31] as described previously [10, 32]. All lifetime histograms used 10 bins per decade. Four different logarithmic probability fits were generated using different fitting algorithms and the mean and standard deviation of the fitted time constants were used as the mean and standard deviation for the characteristic open and blockage times. Power spectrum analysis of the open-channel current noise was performed without digital filtering by ClampFit 10.3. Each experiment was repeated at least three times on different membranes; the data of one experiment are shown for clarity.

### Bilayer Overtone Analysis (BOA) measurements

BOA measurements were performed as described [20, 33] (Supplemental Methods) using a Stanford Research Systems 830 lock-in amplifier and planar lipid membranes made from DOPG/DOPC/DOPE (2:1:1) (mol/mol) in 150 mM KCl buffered with 5 mM HEPES at pH 7.4 by the same protocol as for VDAC reconstitution experiments. The intrinsic membrane potential, Ψ, reports on the asymmetry between two monolayer leaflets. Experimentally, Ψ was determined as described [20, 34] (Supplemental Method). Data are presented as ΔΨ = Ψ — Ψ_*t*=0_, where positive ΔΨ corresponds to the increase of positive charge on the *cis* surface of the membrane. 60 nM αSyn was added to the *cis* compartment leading to increase of ΔΨ; after ΔΨ was stabilized, the aliquots of HK2p in MP solvent were added to both sides of the membrane under constant stirring. Membrane capacitance *C* and Δ Ψ were measured approximately once per minute using in-house Python-based software [34]. Each data point is an average of 10 subsequent ΔΨ measurements taken when the signal reached a steady-state level, as defined by ΔΨ variations within ± 0.5 mV.

### Fluorescence Correlation Spectroscopy (FCS) measurements

Large Unilamellar Vesicles (LUVs) were prepared from DOPG/DOPC/DOPE (2:1:1) (mol/mol) mixture in 150 mM KCl buffered with 5 mM HEPES at pH 7.4 as described previously [20]. Liposomes’ size and polydispersity were determined by light scattering using Zetasizer Nano-ZS90 (Malvern). Homogeneous populations of LUVs of 106 ± 6 nm diameter with a polydispersity < 0.2 were used in measurements. FCS measurements were made in eight-well cover-glass slides (Grace Biolabs) pretreated with Sigmacoat (Sigma Aldrich) before each experiment to prevent liposome and αSyn adhesion to the surfaces as previously described [20], using a Hamamatsu Photonics K.K. C9413-01 spectrometer with a 473 nm excitation laser [35]. Samples for FCS measurements contained 100 nM Alexa488-labeled αSyn alone or in liposome solution (50 mM lipid) in 150 mM KCl, 5 mM HEPES, pH 7.4. with or without HK2p added from its stock solution in MP solvent. To minimize the potential influence of the fluorescent dye on αSyn-membrane binding, Alexa488 was placed in position Y136C in the C-terminus of αSyn.

### Statistics

For the statistical analysis of mean values, the difference between two groups of data were analyzed by a two-tailed *t*-test using *p* < 0.05 as the criterion of significance. Differences between many groups were analyzed by one-way analysis of variance.

## RESULTS

### HK2p selectively targets the mitochondrial outer membrane

We chose the 15 amino acid N-terminal peptide of human hexokinase-2 (HK2p) for proof of principle studies as it is well-known to target mitochondria and may interact directly with VDAC [21, 22, 36, 37]. Using a synthetic construct of HK2p linked to the cell-penetrating peptide (CPP) poly-Arginine (R9) (HK2p-R9) and labeled with tetra-methyl rhodamine dye (TMR) at the C-terminus (HK2p-R9-TMR), we confirmed that HK2p targets mitochondria in HeLa cells. Images in Figure 2A show that the HK2p-R9-TMR construct (red) targets mitochondria (green) in HeLa cells expressing mito-GFP; this is particularly evident in the zoomed-in image in Figure 2A(d). By contrast, a similar construct with a scrambled peptide (ScrP-R9-TMR) displays a diffuse distribution throughout the cytoplasm (Figure 2B). We further confirmed the targeting using N-terminally TMR labeled HK2p-R9 (TMR-HK2p-R9, Supplemental Figure S1) which shows essentially identical targeting behavior to the HK2p-R9-TMR construct compared to diffuse distribution of the scrambled control peptide (TMR-ScrP-R9). Having shown that HK2p selectively targets mitochondria, presumably binding to the outer membrane and/or to VDACs as was suggested previously [21, 22, 36, 37], we tested the peptide’s ability to inhibit αSyn-induced blockage of VDAC reconstituted into the planar membrane.

**Figure 2.**
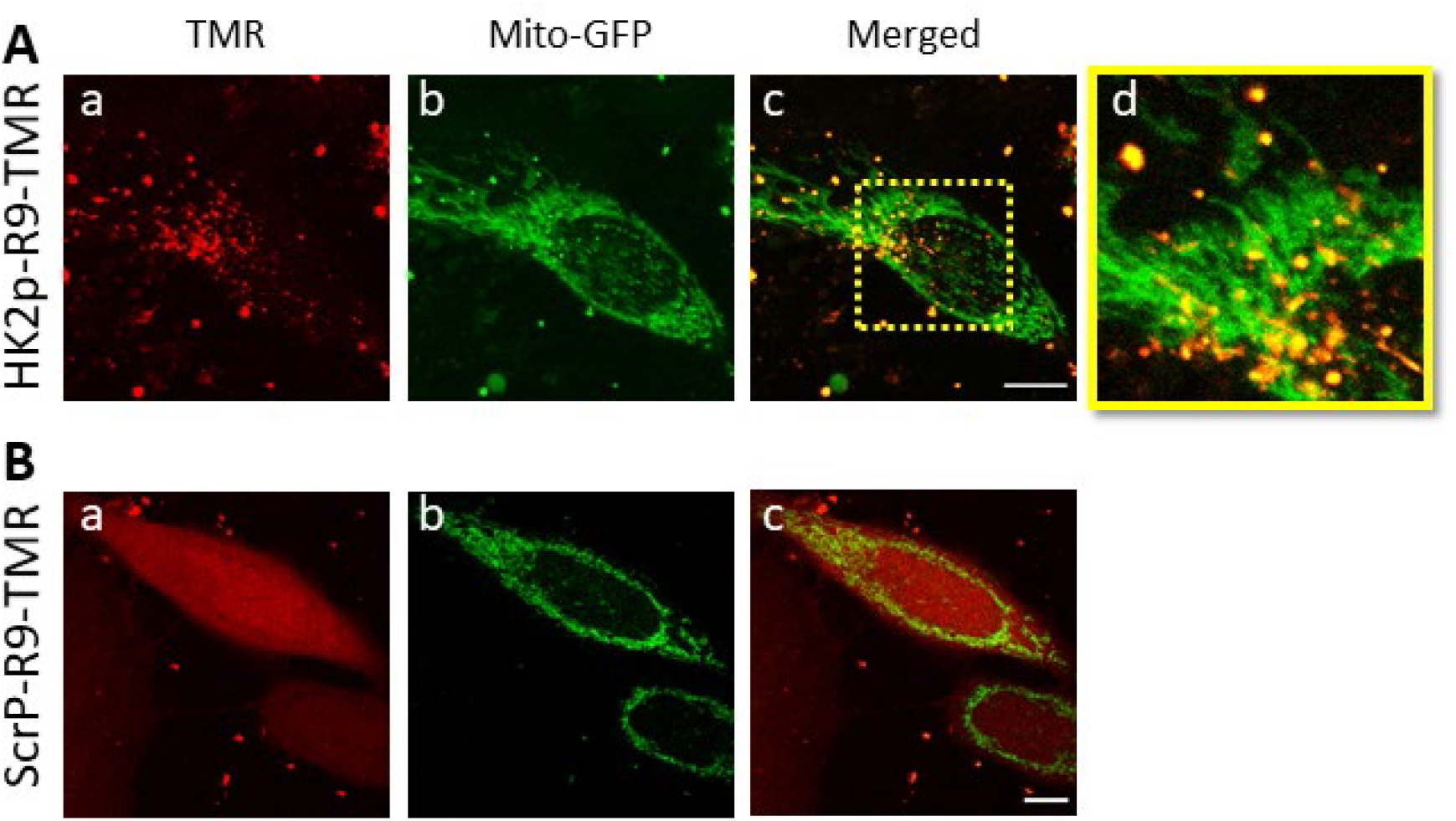
HK2p localization to the mitochondria. (**A**) 5 μM HK2p linked to cell-penetrating peptide poly-Arginine (R9) and tetra-methyl rhodamine dye (TMR), HK2p-R9-TMR (a, red) targets mitochondria in HeLa cells expressing mito-GFP (b, green) as seen by colocalization (c) in the merged image. Zoomed-in image (d) (dashed yellow square in (c)) shows the localization of HK2p-R9-TMR with mitochondria (yellow). (**B**) Control with scrambled peptide (ScrP) shows a diffuse distribution of ScrP-R9-TMR (a) in cells labeled with mito-GFP (b) and does not colocalize with mitochondria (c). A 100x/1.40 oil objective was used for imaging (scale bar 10 μm).

### Membrane-binding peptide inhibits blockage of reconstituted VDAC by αSyn

VDAC reconstituted into a planar lipid membrane (PLM) passes a steady current through the open channel if the applied transmembrane voltage is not sufficiently high to induce gating (Figure 3A, left trace). In this experiment the planar membrane was made from DOPG/DOPC/DOPE (2:1:1) (mol/mol). Addition of αSyn at 50 nM solution concentration induces characteristic two-level fluctuations on the (0.1 to 100) ms time scale, depending on the applied voltage and other experimental conditions (Figure 3A, second from the left trace). The channel conductance fluctuates between that of the open state, which remains the same as in the αSyn-free control, and that of the blocked state, which is ~ 0.4 of the open state conductance. For illustrative clarity, the current traces in Figures 3A were filtered with a digital 1 kHz lowpass Bessel filter, though a larger bandwidth was used for the statistical analysis of current fluctuations. αSyn characteristically blocks VDAC from both sides of the channel, but the blockage events are detectable only when a negative potential is applied to the side of αSyn addition [10], *cis* or *trans* according to the schematic of our PLM system [38]. The situation changes dramatically after the addition of HK2p, in micromolar concentrations, to the same side of the membrane where αSyn was added: the frequency of blockage events decreases progressively with the increase in peptide concentration, with nearly total elimination of the events at 100 μM HK2p (Figure 3A, right traces). The single-molecule nature of these experiments with reconstituted VDAC enables a detailed analysis of the rates of αSyn capture and release by the VDAC pore, *k_on_*, and *k_off_*, respectively [17]. The rates are determined from the distributions of times when the channel is open between successive blockages, *τ_on_*, and is blocked by αSyn, *τ_b_*. The distribution of *τ_on_* is satisfactorily described by a single exponent with the characteristic time *t_on_* = < *τ_on_* > (Figure 3B). The on-rate, which is inversely proportional to *t_on_*, decreases with increasing HK2p concentration (Figure 3C). When normalized to its value in the presence of 50 nM αSyn before HK2p addition, the on-rate can be described as a function of HK2p concentration by a first-order binding equation *k_on_(norm)* = *K_d_*/(*K_d_* +[HK2p]), with *K_d_* = 3.5 ± 0.5 μM. In the representative experiment shown in Figure 3, the hydrophobic HK2p was added to the membrane-bathing solution from its stock dissolved in the mild organic solvent N-Methyl-2-pyrrolidone (MP). A control experiment with an equal volume of MP solvent showed no effect on channel conductance (Supplemental Figure S2A).

**Figure 3.**
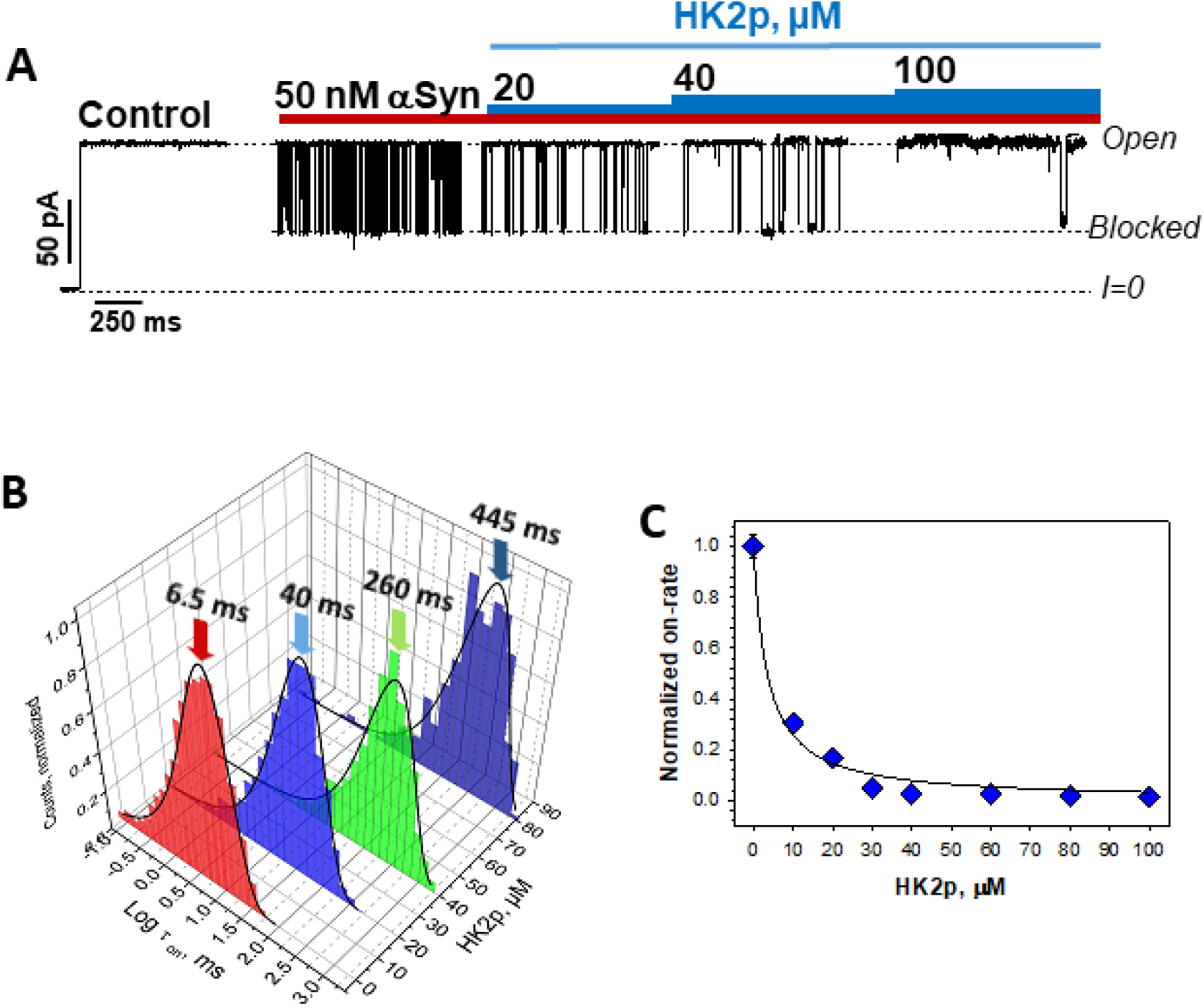
HK2p inhibits αSyn blocking of reconstituted VDAC. (**A**) Representative single-channel experiment showing a current trace through the same VDAC channel before (control) and after *trans* side addition of 50 nM αSyn, followed by sequential addition of 20 μM to 100 μM HK2p to the same side of the membrane, as indicated. Addition of HK2p relieves channel block and promotes the open state. Here and elsewhere horizontal dashed lines indicate VDAC open and blocked states and zero current. For clarity of illustration, the original current record was digitally filtered with a digital 1 kHz lowpass Bessel filter. (**B**) Representative log-binned distributions of open times, *τ_on_*, at different HK2p concentrations. Distributions of *τ_on_* are well described by single-exponential functions (solid black lines), with characteristic times as indicated by arrows. (**C**) The frequency of channel blockages by αSyn (on-rate = 1/<*τ_on_*>, ms^-1^) decreases with HK2p concentration. The normalized on-rate vs HK2p concentration obtained in experiments, an example of which is shown in (A). The on-rate was normalized to the average on-rate in the presence of 50 nM αSyn without HK2p addition. The solid line is the result of a fit to the first order binding equation *k_on_(norm)* = *K_d_*/(*K_d_* +[HK2p]), with *K_d_* = 3.5 ± 0.5 μM. Error bars represent one standard deviation from the mean and are not visible in some as they are smaller than the size of the data point. Planar membrane was formed from DOPG/DOPC/DOPE (2:1:1) (mol/mol). The membrane bathing solution contained 1 M KCl solution buffered with 5 mM HEPES at pH 7.4. Applied voltage was 25 mV. HK2p was dissolved in MP solvent.

To confirm that HK2p, when linked to the CPP, shows similar results, we tested the water-soluble construct of HK2p linked to CPP derived from HIV-1 protein TAT (HK2p-TAT) with VDAC reconstituted into a DPhPC membrane. The TAT peptide itself does not measurably affect αSyn blockage kinetics (Figure 4A). However, addition of the same amount (40 μM) of HK2p-TAT resulted in a significant decrease of the blockage frequency (Figure 4), similar to what was found for the water insoluble HK2p (Figure 3). The average normalized on-rate of VDAC blockages as a function of HK2p-TAT concentration is shown in Figure 4B and is described by a first-order binding curve with *K_d_* = 13.5 ± 1.2 μM. These results confirm that the hydrophobic HK2p domain of the construct is responsible for the reduction of the blockage event frequency and that the presence of the cell-penetrating domain does not qualitatively change this effect.

**Figure 4.**
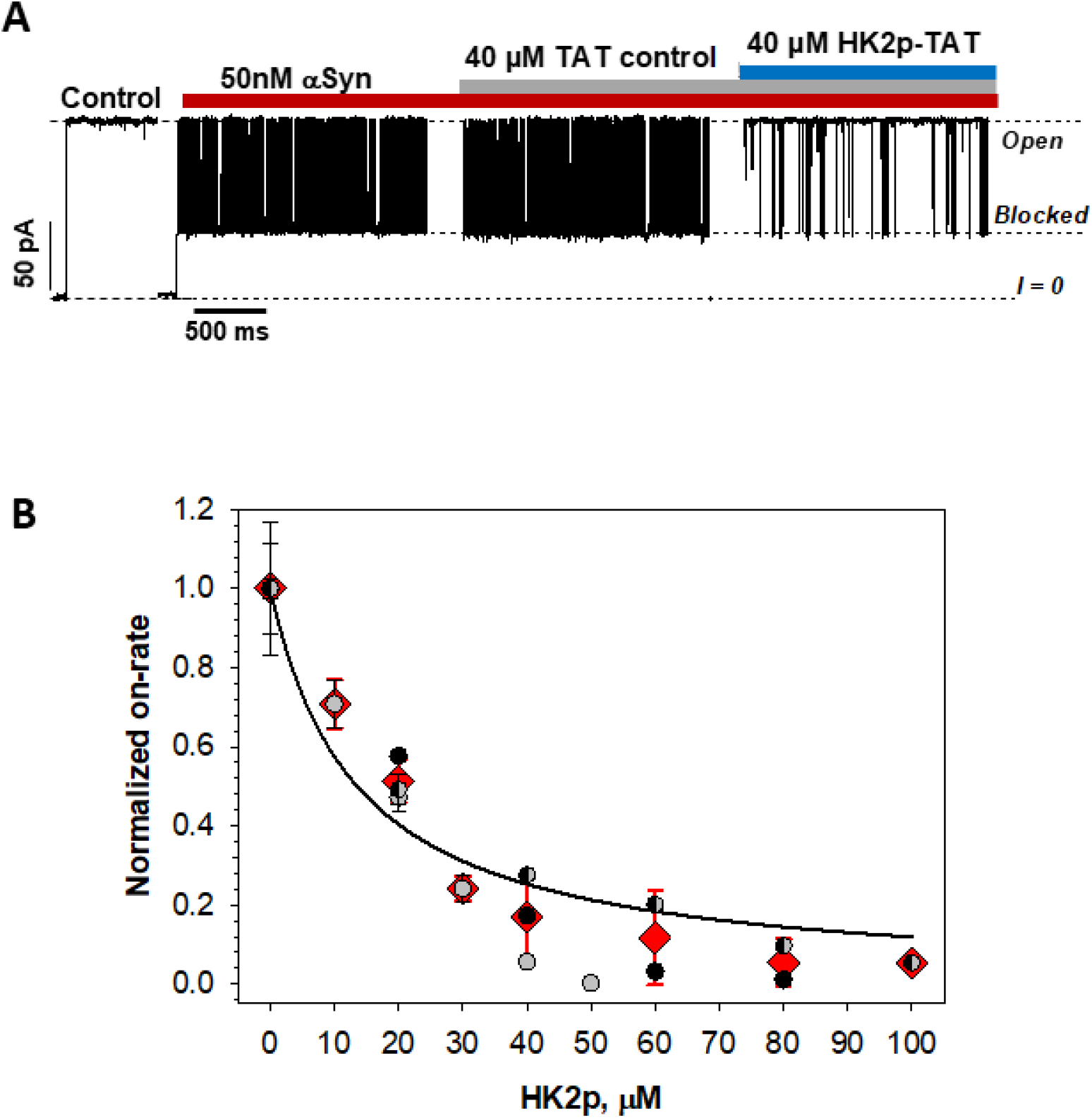
Cell-penetrating HK2p-TAT construct also impedes αSyn interaction with VDAC. (**A**) A representative current trace through the same single VDAC channel reconstituted into DPhPC membrane before (control) and after addition of 50 nM of αSyn (left traces, red bar) followed by addition of 40 μM TAT peptide (TAT control, grey bar) and then 40 μM of HK2p-TAT (blue bar) to the same side of the membrane (*trans* side) as the side of αSyn addition. HK2p-TAT reduces αSyn-induced VDAC blockage events while TAT peptide by itself does not measurably affect the kinetics of VDAC blockage by αSyn. The membrane bathing solution contained 1 M KCl buffered with 5 mM HEPES at pH 7.4. The applied voltage was 30 mV. The original current record was digitally filtered for illustration purposes with a 1 kHz lowpass Bessel filter. (**B**) The normalized on-rate of VDAC blockages as a function of HK2p-TAT concentration. Red diamonds and error bars are the mean and the standard deviation from the mean; other symbols represent data points of the 3 individual experiments. The on-rate was normalized to the average on-rate in the presence of 50 nM αSyn without TAT or HK2p-TAT addition. The solid line is the fit to the simple binding curve with *K_d_* = 13.5 ± 1.2 μM (68% confidence interval estimated from the covariance of the optimized parameter). All experimental conditions as in (A).

The inhibitory effect of HK2p on αSyn-induced blockages was observed with VDAC reconstituted into the planar membranes of different lipid compositions: DOPG/DOPC/DOPE (2:1:1) (mol/mol) (Figure 3), DPhPC (Figure 4), and PLE/Chol/DPhPC (86/9/5) (mol/mol) (Supplemental Figure S2B, C). While DPhPC, though commonly used in channel reconstitution, is not physiologically relevant for the mammalian cells, phosphatidylcholine (PC) and phosphatidylethanolamine (PE) make up 55 and 30%, respectively, of the total content of mammalian MOM [39]. The negatively charged phosphatidylglycerol (PG) used in our lipid mixture represents a high content of anionic lipids in MOM (up to 20%). Therefore, a mixture of DOPG/DOPC/DOPE (2:1:1) (mol/mol) models the polar headgroup composition of a mammalian MOM reasonably well. A mixture of soybean polar lipid extract (PLE) with cholesterol mimics the phospholipid content of rat liver mitochondria even more closely [40] providing a natural mixture of phospholipid acyl chains. Under all studied conditions – whether using the different HK2p constructs or the changed membrane lipid composition – addition of HK2p in ≥ 20 μM concentration resulted in a 10-fold or larger reduction in the on-rate of channel blockage by αSyn, up to a nearly total abolishment of the blockages (as at 100 μM of the HK2p shown in Fig. 3A, C).

By contrast with the open time distributions, which are satisfactorily described with a single exponential function (Figure 3B), the distributions of blockage time, *τ_b_*, become complex with the increasing HK2p concentration and require multi-exponential functions for their description (Figure 5A). At HK2p ≥ 20 μM (a corresponding histogram in Figure 5A is shown in blue), a second population of long-lasting blockage times *τ_off_^(2)^* appears, with the characteristic times ~10 times longer than those of the main component, *τ_off_^(1)^*. Nevertheless, the average blockage time or the αSyn mean residency time in the pore, *t_off_* = <*τ_b_*>, is mostly defined by the main population of the blockage times, *τ_off_^(1)^*, which does not depend on HK2p concentration and remains in the range of 1.3 ms to 1.7 ms (Figure 5A).

**Figure 5.**
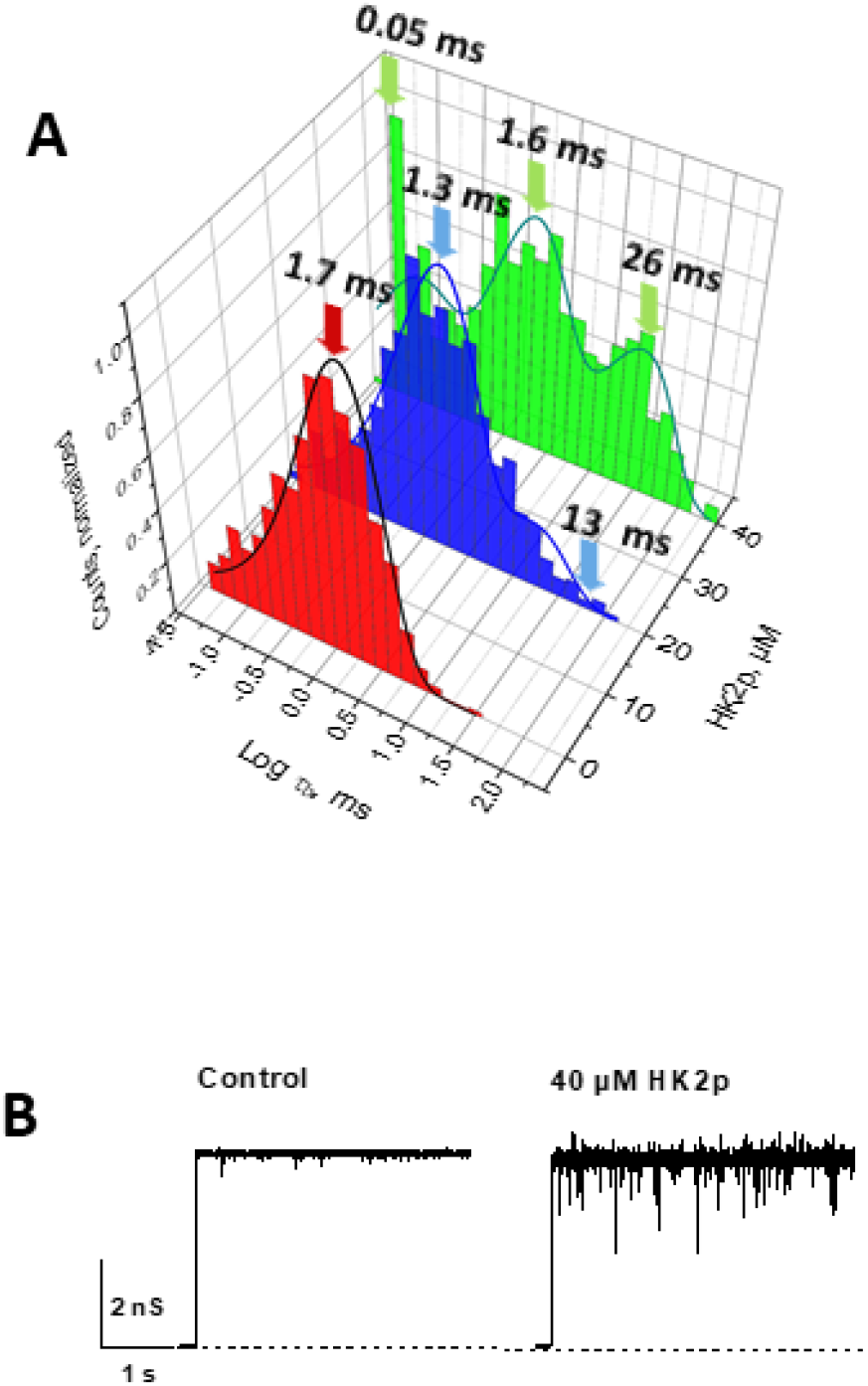
HK2p affects the distribution of αSyn-induced blockage time and induces VDAC open-channel current noise. (**A**) Log-binned histograms of blockage times, *τ_b_* at different HK2p concentrations. The most abundant short-time component (*τ_off_^1^*:1.3 ms to 1.7 ms) does not change appreciably with HK2p concentration. With increasing HK2p concentration (20 μM to 40 μM), a second, longer-lasting component of blockage times appears, and the fitting of *τ_b_* distributions requires two exponential functions (solid lines) with the characteristic times (*τ_off_^1^* and *τ_off_^2^*) indicated. The planar membrane was formed from DOPG/DOPC/DOPE (2:1:1) (mol/mol). Applied voltage was 25 mV. All experimental conditions were as in Figure 3. (**B**) HK2p induces an excess of current noise through the open VDAC state, as witnessed by the representative current records through the same single VDAC channel before and after the addition of 40 μM of HK2p in the absence of αSyn. Channel was reconstituted in DPhPC membrane bathed in 1 M KCl buffered with 5 mM HEPES at pH 7.4. HK2p in MP solvent was added to the *cis* side of the membrane. Applied voltage was −40 mV.

At 40 μM HK2p (the histogram in Figure 5A shown in green), a third, short-lasting component of the blockage times of ~ 0.05 ms is detected. HK2p does not measurably change the conductance of the VDAC open or αSyn-blocked states at the studied peptide concentrations (Figure 3A, 4A, Supplemental Figure S2B). However, at higher HK2p concentrations, a noticeable noise of the current in the open state is observed (Figure 3A, Supplemental Fig. S2B). This phenomenon could be best seen in experiments without αSyn addition which allow recording of relatively long (> 100 s range) current fragments at high applied voltage (typically, the power spectral density of the excess current noise is proportional to the square of the applied voltage [41]). Addition of 40 μM HK2p already induces a measurable open-state current noise independent of αSyn-induced characteristic fluctuations (Figure 5B, Supplemental Figure S3A). An example of the corresponding power spectral density, *S(f)*, of the open channel noise with and without HK2p is shown in Supplemental Figure S3B. The low-frequency spectral density obtained in the presence of 40 μM of HK2p is ~ 15 times higher than in the control and ~ 5 times higher than in the presence of 20 μM of HK2p (Supplemental Figure S3B). The increase in the low-frequency spectral density is due to HK2p-induced fluctuations in the channel conductance and thus indicates a direct peptide-VDAC interaction. These experiments were performed with VDAC reconstituted into DPhPC membranes. To test if the HK2p-induced current noise depends on membrane lipid composition we performed similar experiments with VDAC reconstituted to the PLM made from PLE/cholesterol/DPhPC – and also observed an increase in the open-channel current noise (Supplemental Figure S4A). This suggests that the ability of the HK2p to induce current noise in the VDAC open state does not depend on membrane lipid composition, further supporting the proposed direct interaction between HK2p and VDAC. Control experiments with HK2p solvent MP showed no effect either on the current of the open channel or the unmodified planar membrane (Supplemental Figures S2A, S4B). Therefore, the third, very fast component *τ_off_^(3)^* observed at 40 μM HK2p addition (Figure 5A), most likely represents the current noise induced by HK2p in VDAC open-state current and is not related to the αSyn-induced blockages.

The ability of HK2p to induce noise in the VDAC open-state current (Figure 5B, Supplemental Figure S3) suggests a direct interaction between HK2p and VDAC which was reported previously in a number of studies [25–27]. To further test this possibility, we examined the effect of HK2p on VDAC gating, the characteristic channel property that is sensitive to the changes in experimental conditions such as lipid composition [30, 42, 43], medium pH [44, 45], and osmotic stress [46]. Measurement of VDAC gating allows assessing its interaction with other proteins, low-molecular-weight drugs, and hydrophobic compounds (for a general discussion see [47]) and was shown to be affected by physiologically relevant mitochondrial metabolites such as NADH [48] as well as proteins such as those in the Bcl2 family [49, 50] and cytoskeletal proteins [51, 52]. Thus, changes in VDAC’s gating behavior can be a useful indicator of an interaction with a compound of interest.

The characteristic bell-shaped plots of normalized average VDAC conductance are shown in Supplemental Figure S7A.The addition of 20 μM of HK2p to both sides of the membrane caused a decrease of VDAC gating manifested in increased *G_min_*, the minimum conductance at |*V*| >50 mV. This increase is most pronounced at negative polarities (~ 10% increase in comparison with control) (Supplemental Figure S5A). However, HK2p does not affect gating parameters, *i.e*. the effective gating charge and the voltage at which half of the channels reopen (Supplemental Figure S5B, C). HK2p could modulate VDAC gating by directly interacting with the channel, which goes along with the ability of HK2p to induce current noise of the VDAC open state. Alternatively, HK2p could affect VDAC gating by modifying a surrounding channel lipid (for discussion see [47]).

There are three plausible explanations for how HK2p could counteract the αSyn-induced block of VDAC conductance: (1) hydrophobic HK2p triggers detachment of αSyn from the membrane surface by competing with αSyn for membrane binding sites (a VDAC-independent scenario); (2) HK2p directly interacts with the VDAC protein as was previously suggested [25, 53] and thus competes with αSyn for binding to VDAC (a VDAC-dependent scenario); or (3) a combination of both (1) and (2).

### HK2p competes with αSyn binding to lipid membranes

To test the hypothesis of HK2p-induced depletion of αSyn at the lipid membrane surface, we explored two independent macroscopic methods of measuring αSyn-membrane binding – BOA and FCS.

#### BOA measurements

The second harmonic of the bilayer’s electric response to the applied periodic excitation potential depends on the electrical properties of the bilayer and their modification by any membrane-bound compounds. Specifically, it allows the measurement of a transmembrane potential, *ΔΨ*, caused by the electrical asymmetry between two lipid leaflets. The asymmetry could be induced by different means, including the one-sided addition of a charged peptide that binds to the membrane. The primary advantage of this method for our purposes is that it allows the evaluation of real-time kinetics of binding on the exact same system of PLM as in the VDAC reconstitution experiments [20, 33].

BOA measurements previously showed that αSyn binds to DOPG/DOPC/DOPE (2:1:1) planar membranes [20]. The sign of the resulting ΔΨ asymmetry is positive, confirming the interpretation of different methods that have shown the positively charged N-terminus of αSyn to be the membrane-binding domain [14, 54, 55]. Considering that the *K_d_* of αSyn-membrane binding measured by BOA is ~ 30 nM [20] (which is orders of magnitude lower than the αSyn *K_d_* measured by conventional macroscopic methods; see the relevant discussion in [20]), we used 60 nM of αSyn in BOA competition experiments to ensure a stable binding of αSyn. An example of the time course of transmembrane potential changes obtained on a DOPG/DOPC/DOPE membrane in 150 mM KCl is shown in Figure 6A. The addition of 60 nM αSyn to the *cis* side of the membrane produced a positive ΔΨ ~ 15 mV. The following sequential addition of HK2p symmetrically to both sides of the membrane resulted in a systematic decrease of ΔΨ returning to a baseline of ~ - 2.5 mV after the final addition of 100 μM HK2p (Figure 6A). The binding curve in Figure 6B shows the average ΔΨ as a function of HK2p concentration and demonstrates that HK2p competes with αSyn for membrane binding by effectively displacing it from the membrane with *K_d_* = 26.8 ± 3.0 μM.

**Figure 6.**
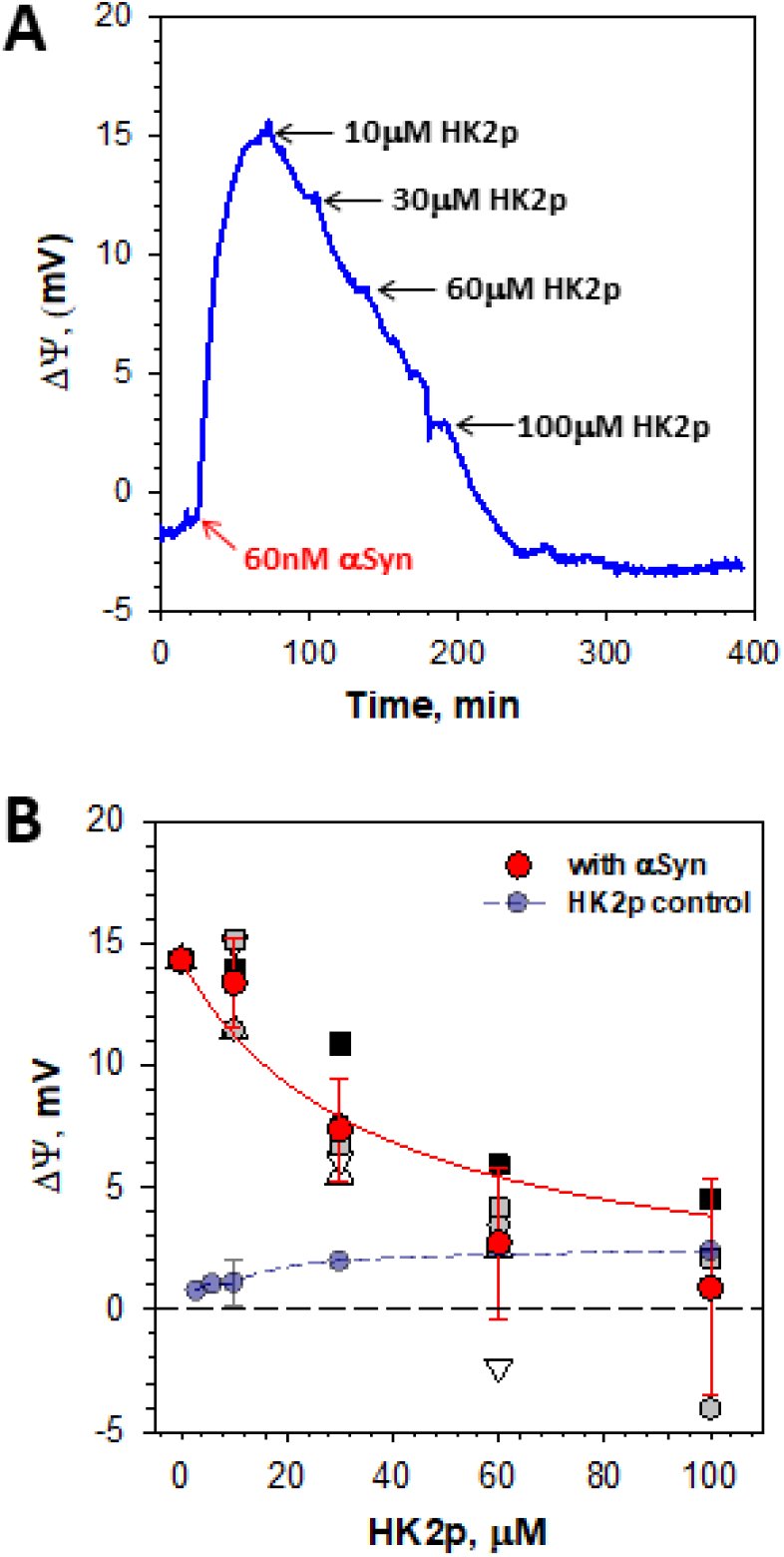
Competition for membrane binding between αSyn and HK2p measured by BOA. (**A**) A representative time course of transmembrane potential (ΔΨ) change due to αSyn binding to the DOPG/DOPC/DOPE (2:1:1) planar membrane in 150 mM KCl, pH 7.4. After stabilization of the ΔΨ ~ 15 mV caused by the addition of 60 nM of αSyn to the *cis* side of the membrane, HK2p was added to both sides of the membrane stepwise in the increasing concentrations indicated by arrows. Each HK2p dose decreases ΔΨ in the direction of the baseline at ΔΨ ~ −2.5 mV. (**B**) Average ΔΨ as a function of HK2p concentration. Red circles and error bars are the mean and standard deviation from the mean; other symbols represent data points of 5 individual experiments. The solid line is a fit to the binding equation with *K_d_* = 26.8 ± 3.0 μM (68% confidence interval estimated from the covariance of the optimized parameter). Control experiment (blue circles) shows that HK2p by itself does not change *ΔΨ* substantially when added to the *cis* side of the membrane.

#### FCS measurements

The BOA method reports exclusively on the electrical asymmetry between the two lipid leaflets. Hypothetically, this asymmetry could be compensated, for example, by the binding of oppositely charged domains of αSyn to the membrane upon interaction with HK2p. To exclude such a possibility, we performed FCS measurements that report on αSyn dissociation constants independently of the charge of its membrane binding domains [20, 56, 57]. The FCS normalized autocorrelation curves (AC) for free Alexa-488-labeled αSyn and in the presence of Large Unilamellar Vesicles (LUVs) made of the same lipid composition as PLM in BOA experiments (DOPG/DOPC/DOPE 2:1:1) in 150 mM KCl are shown in Figure 7 (blue and red symbols, respectively). The addition of liposomes reduces the number of free labeled αSyn decreasing their contribution to the AC due to αSyn-liposome membrane binding previously shown for various lipid compositions [20, 56, 57]. This effect results in the shift in the autocorrelation characteristic time towards longer times, which could be seen by comparison of the normalized ACs of the free labeled αSyn and in solution with LUVs (indicated by orange arrow in Fig. 7). The normalized AC shows a substantial shift in the autocorrelation characteristic time towards shorter times with HK2p addition (Figure 7, gray arrow), which represents a decrease of the fraction of LUV-bound αSyn and a corresponding increase of free αSyn. Thus, the HK2p-induced detachment of labeled αSyn from liposome membranes can be seen in the raw AC curves.

**Figure 7.**
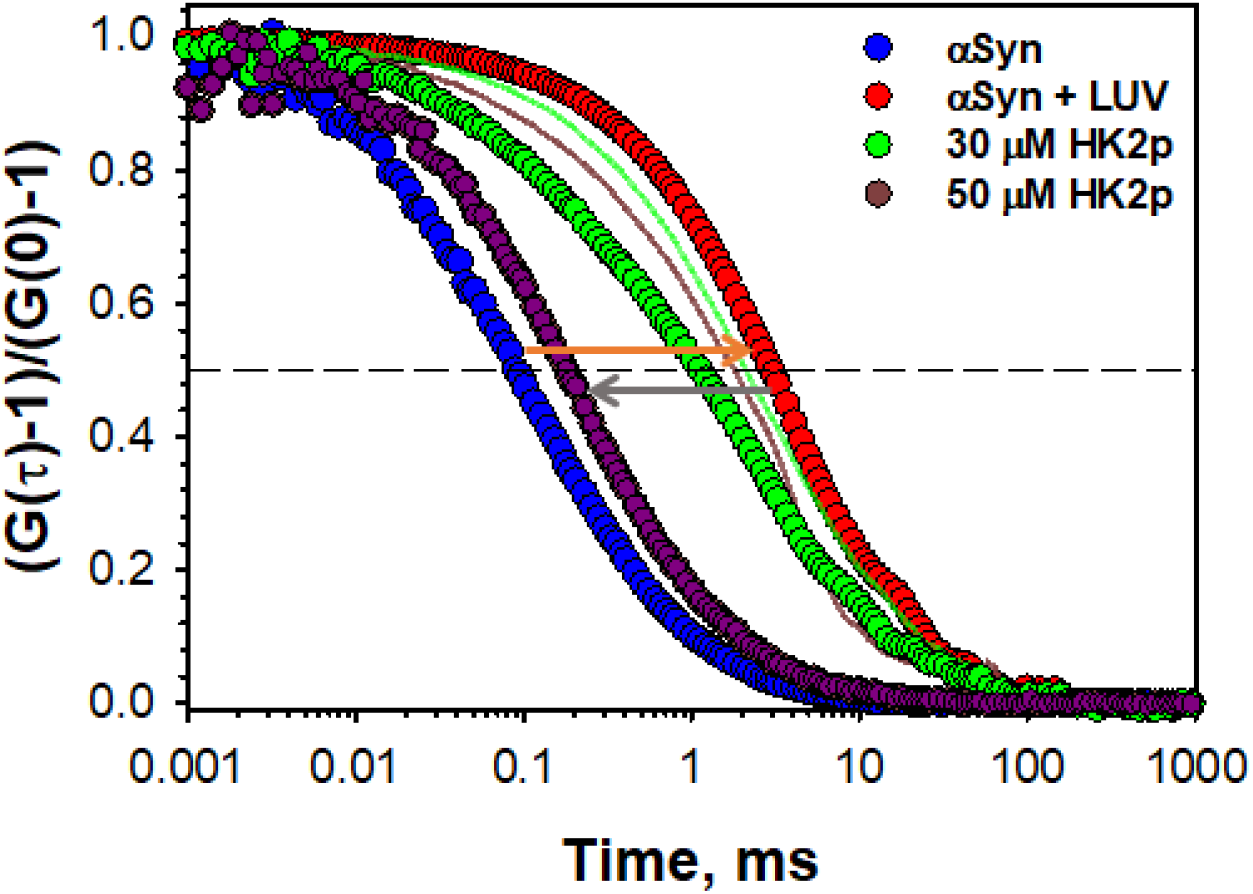
HK2p induces detachment of αSyn bound to liposome membranes measured by FCS. Representative normalized autocorrelation curves show an example of the increase in characteristic autocorrelation time (indicated by the orange arrow) from free unbound αSyn (blue circles) to liposomes-bound αSyn (red circles). The addition of HK2p to the mixture of αSyn and LUVs results in a time shift (indicated by gray arrow) of the autocorrelation curves for 30 μM (green circles) and 50 μM (purple circles) of HK2P concentrations towards free unbound αSyn. The solid lines are the control autocorrelation curves obtained in the presence of aliquots of the HK2p solvent MP corresponding to 30 μM (green line) and 50 μM (brown line) of HK2p addition, respectively. The autocorrelation signal, (*G(t)* – 1), is normalized over the corresponding amplitude of the signal at *τ* = 0.001 ms, (*G(0)* – 1). Liposomes were formed from DOPC/DOPE/DOPG (1:1:2) (mol/mol) mixture (50 μM lipid) in 150 mM KCl buffered by 5 mM HEPES at pH 7.4. The concentration of Alexa-488-labeled αSyn was 100 nM.

The FCS results show that HK2p induces depletion of labeled αSyn on liposome membranes, which is consistent with our interpretation of BOA results. This effect of HK2p on αSyn membrane binding found independently of the VDAC presence could explain the inhibitory effect of HK2p on the αSyn-VDAC complexation. If this model of the inhibition is correct, then one can argue that HK2p should also reduce the αSyn blockage of other β-barrel nanopores, unrelated to mitochondria or VDAC.

### HK2p reduces αSyn blockage of the α-hemolysin β-barrel channel

To test the hypothesis that the effect of HK2p on the αSyn-VDAC complexation does not depend on specific interactions between αSyn and VDAC, we performed experiments with perhaps the most extensively studied biological nanopore, α-hemolysin (αHL) [58–60]. αHL resembles VDAC only in that it is also a β-barrel channel; otherwise, there are few similarities (see discussion in [17]). The constriction of αHL is long, nearly 5 nm, and is formed by a heptameric β-barrel [61]; VDAC is monomeric with a shorter constriction formed by an α-helix lying along one side of the β-barrel [62]. αHL is pronouncedly asymmetric, with a large vestibule on the membrane *cis* side; VDAC has no extramembrane domains. Finally, αHL is essentially nonselective, compared to the substantial anion selectivity of the VDAC channel. Despite these differences, Gurnev et al. showed that nanomolar αSyn added to the stem side of αHL (Supplemental Figure S6A) induces characteristic blockage events of channel conductance with duration in the range of 1 ms to 1000 ms [58]. As in the experiments with VDAC, the blockages of αHL are observed only at negative voltages applied at the side of αSyn addition [58]; a significant rate of such captures is only observed from the side of the membrane opposite the αHL vestibule (Supplemental Figure S6A). In the present study, we found that 20 μM of HK2p added to the same side of the PLM as 50 nM of αSyn caused a pronounced reduction of the rate of αHL blockage events (Supplemental Figure S6B). Thus, HK2p noticeably reduces the blockage of αHL at the same concentration range as was found for VDAC, confirming the proposed mechanism of nanopore-independent competition between HK2p and αSyn on the membrane surface. These results also provide additional support to the mechanism of the αSyn-VDAC complexation wherein the αSyn-membrane binding step precedes αSyn capture by the nanopore.

### HK2p reduces αSyn entry into mitochondria

As was previously shown by different approaches using reconstituted VDAC and live cells, αSyn not only reversibly blocks the VDAC pore but can translocate through VDAC into mitochondria [10, 11, 19]. Therefore, if HK2p reduces VDAC blockage by αSyn *in vitro*, it is expected to also reduce αSyn entering mitochondria in live cells. To test this hypothesis, we used *in situ* PLA [63] which has been proven to be a valuable method to visualize the co-localization of αSyn, mainly a cytosolic protein, and mitochondrial membrane proteins [8, 11]. PLA dots (Figure 8A) represent proximity between αSyn and COX IV at the IM in HeLa cells expressing endogenous αSyn and thus indicate the presence of αSyn in the intermembrane space. We confirmed the localization of PLA dots at mitochondria in untreated cells using Alexa fluor-488 labeled antibody against Tom20, a MOM protein used as a reliable mitochondrial marker (Figure 8A, merged). The PLA time course shows a gradual decrease of PLA signal (number of dots per cell) (Figure 8B) with time and a significant decrease after 10 min (*p*<0.05) incubation with 5 μM HK2p (Figure 8C and Supplemental Figure S7A). Treatment of cells with 5 μM HK2p for 10 mins resulted in a significant reduction (*p*<0.05) of the PLA signal compared to treatment with 5 μM ScrP control (Figure 8B, D, and Supplemental Figure 7B). Considering that VDAC is the main pathway for αSyn entering mitochondria, these results confirm the inhibitory effect of HK2p on αSyn translocation into mitochondria through VDAC in live cells.

**Figure 8.**
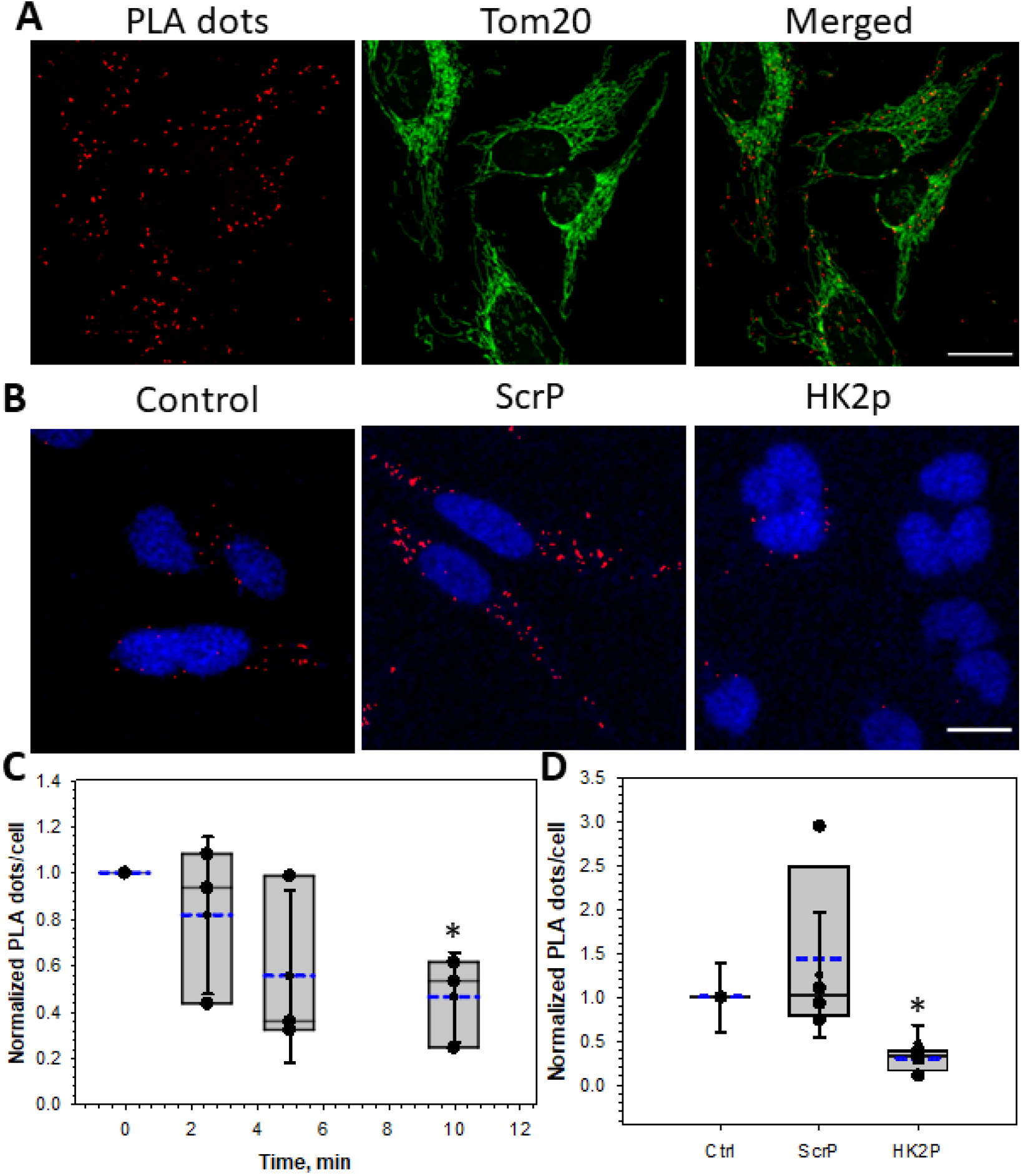
HK2p reduces αSyn entry into mitochondria. (**A**) Representative image shows that PLA between αSyn and COX IV (red dots) colocalize (merge) with mitochondrial network labeled with Tom20-Alexafluor488 (green) imaged with 100x/1.40 oil objective (scale bar 20 μm). (**B**) Representative images showing a decrease in PLA dots after the addition of 5 μM HK2p compared to the addition of 5 μM scrambled peptide (ScrP). The nuclei are stained with DAPI (blue) and imaged with 20x/0.75 objective (scale bar 20 μm). (**C**) Box plots represent the normalized PLA signal per cell (normalized to time 0 without HK2P addition) at different times after the addition of 5 μM HK2p, obtained from an average of 65 cells per experiment with a minimum of 30 cells in 3 independent experiments. (**D**) Box plots represent the PLA dots per cell upon addition of 5 μM ScrP vs HK2P normalized to control (Ctrl) without any treatment, obtained from an average of 23 cells per experiment with a minimum of 5-10 cells in 4 independent experiments. The borders of the boxes define the 25^th^ and 75^th^ percentiles, with the dashed blue line as a mean, the median displayed as black lines, and error bars indicating standard deviation from the mean. Significance was tested against the control without treatment using a paired t-test (* *p* ≤ 0.05).

## DISCUSSION

Here, as a proof of principle, we demonstrated the inhibitory effect of a synthetic mitochondria membrane-binding peptide on the αSyn complexation with VDAC. The obtained results also confirm our previous discovery that αSyn binding to the lipid membrane is the first and crucial step in the complex formation [17, 19, 20] (Figure 1). Single-channel experiments with VDAC show that HK2p, added in micromolar concentrations to the same side of the membrane as αSyn, considerably reduces the number of VDAC blockage events caused by αSyn and, at 60 μM to 100 μM, essentially eliminates the blockage (Figures 3, 4). Notably, this inhibitory effect of HK2p was observed on VDAC reconstituted into the membranes made of different lipid compositions (Figures 3 and 4 and Supplemental Figure S2). Competition experiments performed by independent BOA and FCS methods, using planar and liposomal membranes, respectively, show that HK2p induces detachment of αSyn from the membrane surface with *Kd* ~ 30 μM (as calculated from BOA experiments) and that this process is VDAC-independent (Figures 6, 7). The proposed model of αSyn-VDAC interaction is rather general (Figure 1) because it does not include explicit constraints for either nanopore geometry or specific binding sites, as long as the pore satisfies the requirement of being large enough to accommodate a disordered polypeptide chain. Indeed, the single-channel experiments with αHL confirm this conjecture. They also demonstrate that HK2p effectively reduces αSyn blockages of αHL at solution concentrations similar to those required for VDAC (Supplemental Figure S6).

Recently, Haloi and coauthors [25] studied the complex formation between hexokinase-2 and VDAC using an *in silico* approach. They found that this is a two-step process, where the first step is the membrane binding of hexokinase-2, and the second step is its direct interaction with VDAC. Their molecular dynamics simulations suggested that hexokinase-2 binds to the membrane by inserting its N-terminal domain (or HK2p) into the bilayer independently of VDAC. Using Brownian Dynamics (BD) simulations, they found that the membrane-bound hexokinase-2 could interact with VDAC1 by forming a complex between its membrane-inserted N-terminal domain and the VDAC-lipid interface. These computational results demonstrate the possibility of a direct interaction between HK2p and VDAC protein. Our findings showing the effect of HK2p on the open-channel current noise (Figure 5B and Supplemental Figure S3) and VDAC gating (Supplemental Figure S5) also point to a direct HK2p-VDAC interaction and thus support the model [25]. The negligible effect of HK2p on the average conductance of VDAC in the open and αSyn blocked states and on the dynamics of αSyn escape from the VDAC pore (average blockage time, Figure 5A) suggests that the hydrophobic peptide indeed interacts with VDAC at the protein-lipid interface and, potentially, with VDAC’s external loops as proposed by Haloi and coauthors [25]. Though we cannot exclude an effect of direct HK2p-VDAC interactions on αSyn-VDAC complexation, all data presented here—from BOA and FCS measurements and especially from the independent experiments with αHL—point towards the conclusion that the primary mechanism of HK2p’s inhibitory effect is competition with αSyn for membrane binding sites. Therefore, this is the primary mechanism and sufficient to reduce αSyn-VDAC complexation (Figure 1).

The complex effect of HK2p on the distributions of VDAC blockage time (Figure 5A) yields important insight into the nature of the HK2p-induced inhibition of αSyn membrane binding. At concentrations of 20 μM and above, HK2p causes the emergence of a second, long-lasting component of blockage time, *τ_off_^(2)^*, while retaining a substantial population of events characterized by the average blockage time in the absence of HK2p, *τ_off_^(1)^*. We have previously described a similar phenomenon for VDAC reconstituted into membranes of different lipid compositions [19]. Using energy landscape modeling, we demonstrated that the complex *τ_off_* behavior can be understood in terms of the membrane binding energy and the penetration depth of the anionic CTT of αSyn into the VDAC pore [17]. In particular, the population of the long-lasting *τ_off_^(2)^* is characterized by a much longer penetration depth than that of the short-lived *τ_off_^(1)^*. This phenomenon fundamentally arises from the multiplicity of residues and lipids involved in the binding of a single αSyn molecule. When bound to neutral or lamellar lipids, αSyn is unstructured [16, 64, 65] and can thus bind in an ensemble of conformations. Some of these conformations involve residue-lipid interactions near the CTT [55] so that the entire anionic CTT domain cannot traverse the pore before being arrested by the membrane-bound domain. The polyanionic CTT is the region of αSyn in the nanopore at this point, and the electric field in the nanopore thus destabilizes the membrane binding and favors translocation [17]. These conformations are responsible for *τ_off_^(1)^* and are the only conformations observed on anionic lipids, which include residue-lipid interactions up to residue 94 of αSyn [66]. On the other hand, some conformations do not involve residue-lipid interactions near the CTT. The entire CTT and some of the N-terminal domain is free to traverse the pore, so that the sparsely charged N-terminal domain is more likely to experience the electric field in the channel. This reduces the average electrical forces acting on the polypeptide strand, thus increasing the αSyn-VDAC complex stability and, importantly, disfavoring translocation. These conformations comprise events characterized by *τ_off_^(2)^* [17]. The emergence of *τ_off_^(2)^* is thus the result of loose, unstructured conformations of αSyn on the membranes made of the lipids with low αSyn affinity [17, 20].

By contrast, the experiments presented here have been performed on membranes made of lipids with high αSyn affinity—DOPG/DOPC/DOPE and DPhPC [20, 57]—and thus with one characteristic blockage time [10, 19]. Indeed, these two lipid compositions were chosen intentionally with the aim of avoiding the known complexity of *τ_b_* distributions. The appearance of *τ_off_^(2)^* in the presence of HK2p, therefore, suggests that HK2p disrupts the preferred binding conformation of αSyn molecules, either by modifying the membrane domains to which αSyn binds or by directly interacting with αSyn on the membrane surface, resulting in the detachment of the membrane binding domain near the CTT. It is also possible that HK2p occupies the surface with sufficiently high density so that not all αSyn molecules can adopt their preferred, all-helical conformation. The persistence of *τ_off_^(1)^* simply suggests that not all αSyn molecules are affected. Furthermore, it was shown that events characterized by *τ_off_^(2)^* mostly do not correspond to αSyn translocation, contrary to those characterized by *τ_off_^(1)^*[19]. We thus anticipate an even stronger reduction αSyn translocation through VDAC in the presence of HK2p, compared with that caused by the mere decrease of the on-rate.

The inhibitory effect of HK2p on αSyn entering mitochondria demonstrated on live cells supports the *in vitro* results and can be explained by the proposed mechanism. The toxic effect of monomeric (and oligomeric) αSyn on mitochondrial function is quite complex and involves the interaction of αSyn with a panoply of mitochondrial proteins, such as the TOM complex on the MOM and the ETC complexes on the inner membrane, as well as effects on Ca^2+^ homeostasis, permeability transition, and mitochondrial fission [7, 8, 67–70]. Our results suggest that peptide inhibitors may effectively reduce mitochondrial dysfunction in PD by reducing αSyn in mitochondria (Figure 8). Even though HK2p is known to induce apoptosis by causing detachment of hexokinase-2 from mitochondria in cancer cells where it is overexpressed [21–23, 36, 37], the cells were healthy in our experiments with ≤ 5 μM of HK2p-CPP for at least 10 mins after HK2p addition. Further toxicity studies in neurons will be needed to explore the potential of this peptide for PD treatment.

We believe that the HK2p-CPP construct could be used in the future as a template for creating potent peptide therapeutics against αSyn toxicity which should satisfy four requirements: (1) selectively target the MOM; (2) bind to the membrane, (3) be cell-permeable, and (4) be non-toxic. The type of membrane-binding peptide is not otherwise expected to be important, provided it competes effectively with αSyn for membrane binding sites and preferably at even lower concentrations. This study proves that the search for more potent peptides that are able to reduce αSyn recruitment to MOM, and thus its complexation with VDAC and perhaps with some other mitochondrial proteins, is likely to be a promising endeavor. We expect this type of peptide to play a neuroprotective role by inhibiting αSyn complexation with VDAC and consequently promoting efficient ATP/ADP exchange across MOM while decreasing calcium uptake. Further inhibition of αSyn translocation into mitochondria through VDAC should decrease ROS production and restore mitochondrial potential and oxidative phosphorylation (Figure 9). We believe that VDAC represents an important therapeutic target in neurodegeneration and that compounds that disrupt VDAC interaction with neuronal toxins could provide regenerative or protective therapies against neuronal death in PD.

**Figure 9.**
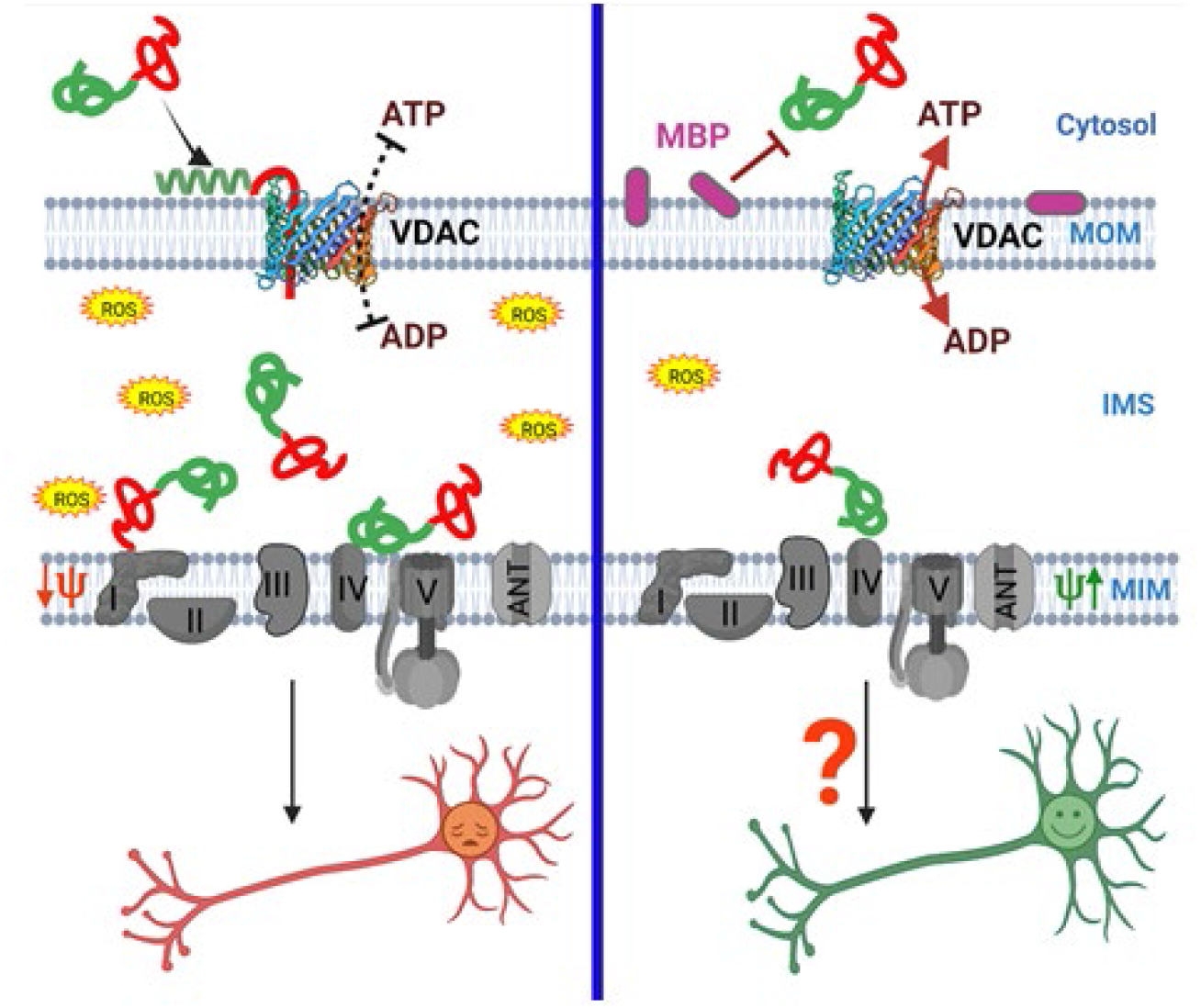
Proposed model of neuroprotection by the mitochondrial outer membrane (MOM)-binding peptide (MBP). αSyn complexation with VDAC results in decreased ATP/ADP exchange through the channel and entering mitochondrial intermembrane space (IMS) through VDAC, αSyn targets ETC complexes in the inner membrane (IM) causing their impairment and leading to the loss of mitochondrial potential (ψ_im_), and increase of reactive oxygen species (ROS) production (*left panel*). The neuroprotective role of MBP is achieved by inhibition of αSyn complexation with and translocation through VDAC. MBP induces detachment of αSyn from MOM thus promoting VDAC open state and efficient ATP/ADP exchange to restore ψ_im_ and enhance oxidative phosphorylation (*right panel*). Created with BioRender.com.

## Authors Contribution

Tatiana Rostovtseva, Megha Rajendran, David Hoogerheide, María Queralt-Martín, Philip Gurnev, and Sergey Bezrukov contributed to the study conception, experimental design, and methodology. Material preparation, data collection, and analysis were performed by Megha Rajendran, María Queralt-Martín, Philip Gurnev, William Rosencrans, Amandine Rovini, Daniel Jacobs, Kaitlin Abrantes. The first draft of the manuscript was written by Tatiana Rostovtseva and Megha Rajendran. Review and editing of the manuscript were performed by Tatiana Rostovtseva, Megha Rajendran, David Hoogerheide, María Queralt-Martín, Amandine Rovini, William Rosencrans, and Sergey Bezrukov. All authors read and approved the final manuscript.

## Acknowledgements

This work was started by and is dedicated to the memory of Philip Gurnev. M.R., M.Q.M, P.A.G., W.M.R., D.J., A.R., K.A., S.M.B., and T.K.R. were supported by the Intramural Research Program of the National Institutes of Health (NIH), Eunice Kennedy Shriver National Institute of Child Health and Human Development. M.Q.M also acknowledges support since 2020 from postdoctoral fellowship Juan de la Cierva Incorporación from the Spanish Ministry of Science and Innovation MCIN/AEI/ 10.13039/501100011033 (IJC2018-035283-I, 2020-2022) and support since 2021 from Universitat Jaume I (project UJI-A2020-21). Certain commercial materials, equipment, and instruments are identified in this work to describe the experimental procedure as completely as possible. In no case does such an identification imply a recommendation or endorsement by the National Institute of Standards and Technology (NIST), nor does it imply that the materials, equipment, or instruments identified are necessarily the best available for the purpose.

