## Supplemental Information for "Restricting α-Synuclein Transport into Mitochondria by Inhibition of α-Synuclein-VDAC Complexation as a Potential Therapeutic Target for Parkinson’s Disease Treatment"

### SUPPLEMENTAL METHODS

#### BOA measurements and data analysis

The BOA measurements were performed as described [1, 2] using a Stanford Research Systems 830 lock-in amplifier and planar lipid membranes made from DOPG/DOPC/DOPE (2:1:1) (mol/mol) in 150 mM KCl buffered with 5 mM HEPES at pH 7.4. An excitation potential *V(t)* with frequency $f_0 = 1933\ Hz$ and amplitude $V_{ac} = 106\ mV\ (75\ mV_{rms})$, and variable dc potential $V_{dc}$ were applied to the membrane. The ac current was measured at the frequency of the second harmonic $2f_0 = 3866\ Hz$. For a lipid bilayer membrane with a capacitance *C* that scales with applied voltage as $C = C_0 + \alpha V^2$, where $C_0$ is the capacitance in the absence of an applied potential and $\alpha$ is related to the compressibility of lipid bilayer, the second harmonic current is [1, 3]:

$$i_2(t) = -6\pi f_0 \alpha (\Psi + V_{dc}) V_{ac}^2 \sin(4\pi f_0 t). \quad (1)$$

Here Ψ, the intrinsic membrane potential, reports on the asymmetry between two monolayer leaflets. Experimentally, Ψ was determined as described [1, 2]: from measuring $i_2(t)$ amplitude as a function of $V_{dc}$, which was swept from –50 to 50 mV in steps of 10 mV; the amplitude is minimized at $V_{dc} = -\Psi$ .

### VDAC gating experiments

VDAC's voltage-dependent properties were assessed following the protocol previously devised [4-6] where gating is inferred from the channels response to a slow symmetrical 5 mHz triangular voltage wave of ± 60 mV amplitude from an Arbitrary Waveform Generator 33220A (Agilent). Data were acquired at a sampling frequency of 2 Hz using pClamp 10.7 software and analyzed following published protocols [4, 5] using an algorithm developed in-house. In each experiment, current records were collected from membranes containing 10-100 channels in response to 5-10 periods of voltage waves, to ensure a minimum of 100 channels per experiment. Only the part of the wave during which the channels were reopening was used for the subsequent analysis [7].

Analysis of VDAC voltage gating was performed following previously described protocols [4, 5]. Given the variable number of channels during the experiment, the average conductance ($G$) was normalized to the maximum conductance ($G_{max}$). The probability of the channel to be open, $P_{open}$, is defined as:

$$P_{open} = \frac{G - G_{min}}{G_{max} - G_{min}}, \qquad (2)$$

where $G_{max}$ and $G_{min}$ are the maximum and minimum conductances, corresponding to the channels mostly open at small voltages (<10 mV) and the channels mostly closed at high voltages (>30 mV), respectively. $P_{open}$ plots were fit according to the Boltzmann equation:

$$P_{open} = \left(1 + \exp\left(\frac{F}{RT} \cdot n(|V| - V_0)\right)\right)^{-1}, \qquad (3)$$

where $V_0$ is the voltage at which half of the channels are open, $n$ is the effective gating charge, and $R$, $T$, and $F$ are the gas constant, absolute temperature, and Faraday constant, respectively.

**SUPPLEMENTAL FIGURES**

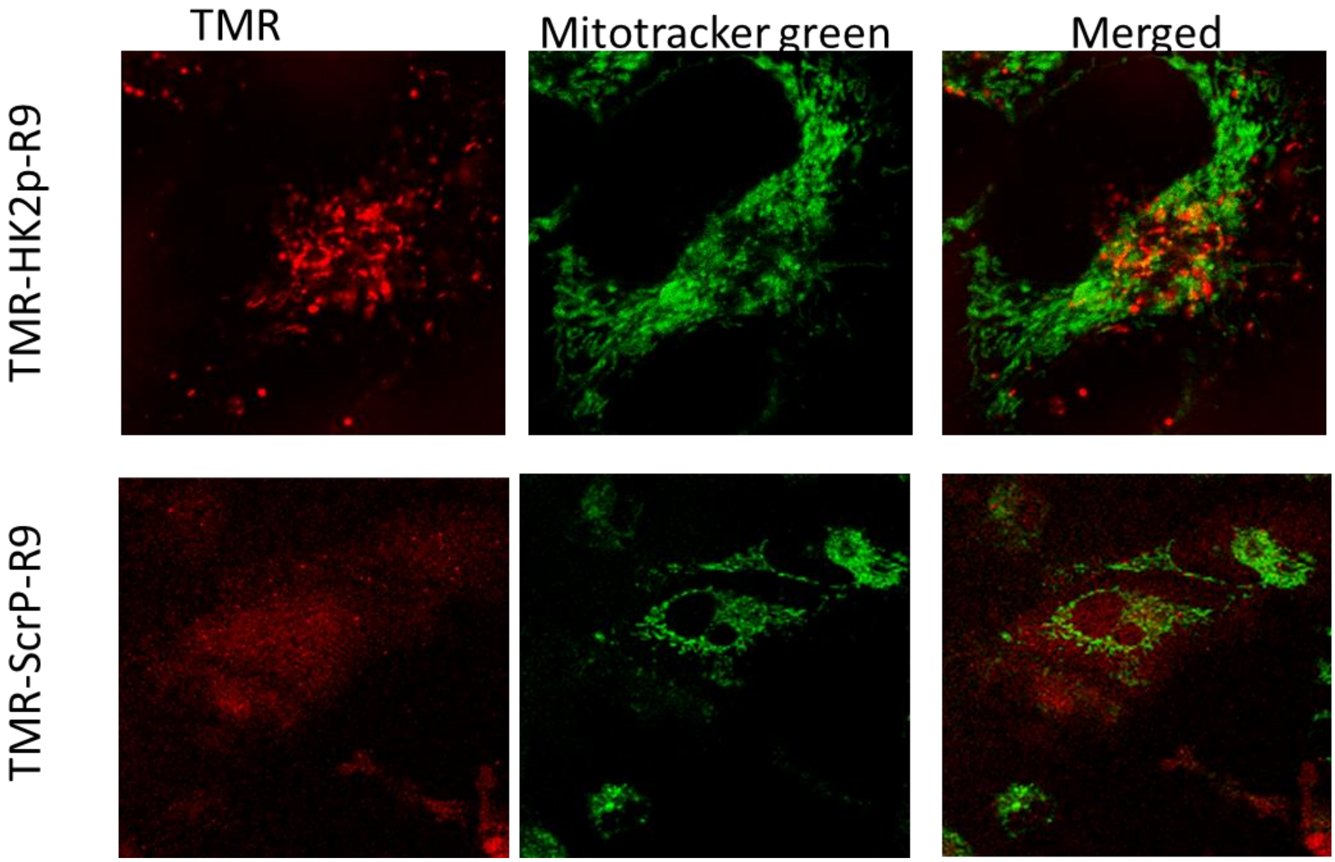


**Supplemental Figure S1.** HK2p localization to the mitochondria. 5 µM HK2p linked at C-terminal to cell-penetrating peptide poly-Arginine (R9) and N-terminal labeled with tetramethyl rhodamine (TMR), TMR-HK2p-R9 (red), targets to the mitochondria (green) in HeLa cells stained with mitotracker green as seen by colocalization (yellow) in the merged image. A similar construct with Scrambled peptide (TMR-ScrP-R9) has a diffuse distribution in cells.

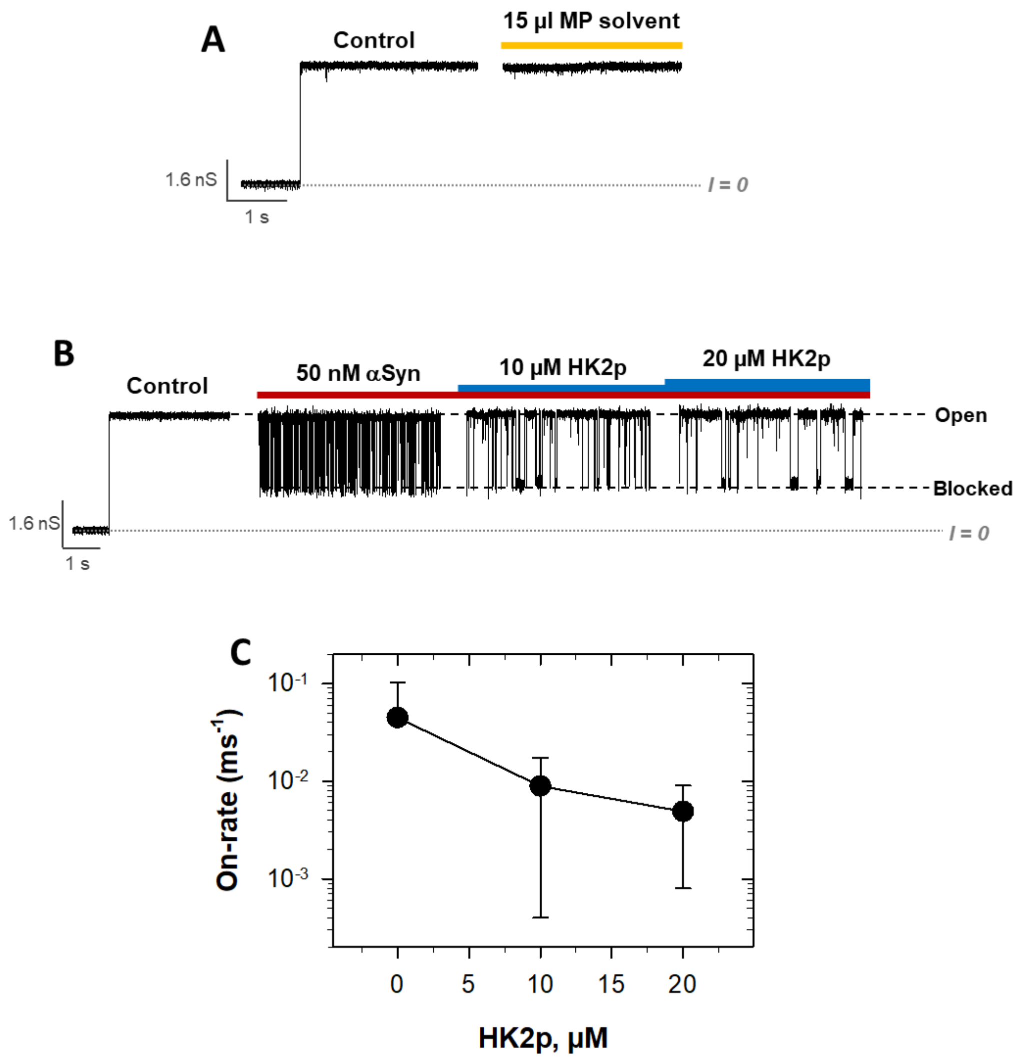


**Supplemental Figure S2**. HK2p reduces αSyn block of VDAC reconstituted in PLE/cholesterol membranes. (**A**) Control single-channel experiment showing the absence of the effect of MP solvent of VDAC conductance. 15 µl of MP solvent would correspond to an addition of 20 µM HK2p. (**B**) Representative single-channel experiment showing a current trace through the VDAC channel at -25 mV applied voltage, before (control) and after addition of 50 nM αSyn, followed by sequential addition of HK2p in MP solvent at 10 µM and 20 µM, as indicated. Subsequent addition of HK2p relieves channel block and promotes the open VDAC conductance. Planar membrane was formed from PLE/Chol/DPhPC (86/9/5) (mol/mol). The membrane bathing solution contained 1 M KCl buffered with 5 mM HEPES at pH 7.4. αSyn and HK2p were added to the *cis* side of the membrane. Horizontal dash lines indicate VDAC open and blocked states and a dotted line indicates zero current. The original current record was digitally resampled-averaged with 10 as a smoothing factor. (**C**) Averaged on-rate at different HK2p concentrations. Data points and error bars represent the mean and its 68% confidence interval, as estimated from the standard deviation from the mean from two independent experiments.

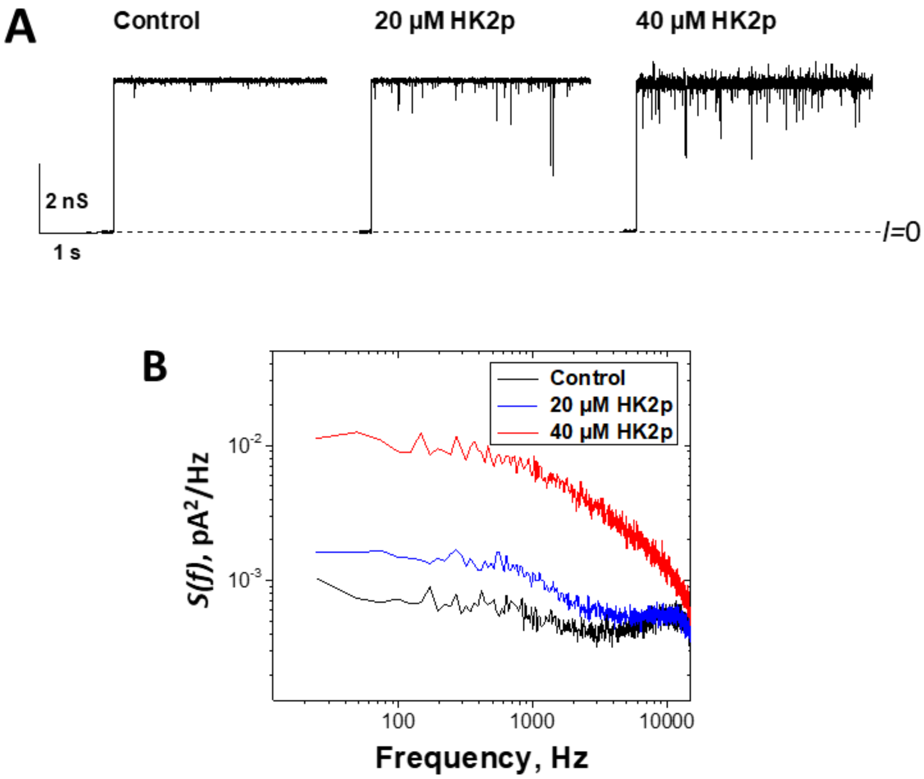


**Supplemental Figure S3**. HK2p induces an excess of current noise through the open VDAC state. (**A**) Representative current records through the same single VDAC channel before and after the addition of 20 and 40 µM of HK2p. Consecutive additions of HK2p induce an increase in the open-channel current noise. Dotted line indicates zero current. (**B**) Power spectral density of current noise through a single fully open channel increases with HK2p addition. Channel was reconstituted in DPhPC membrane bathed in 1 M KCl buffered with 5 mM HEPES at pH 7.4. HK2p in MP solvent was added to the *cis* side of the membrane. Applied voltage was -40 mV.

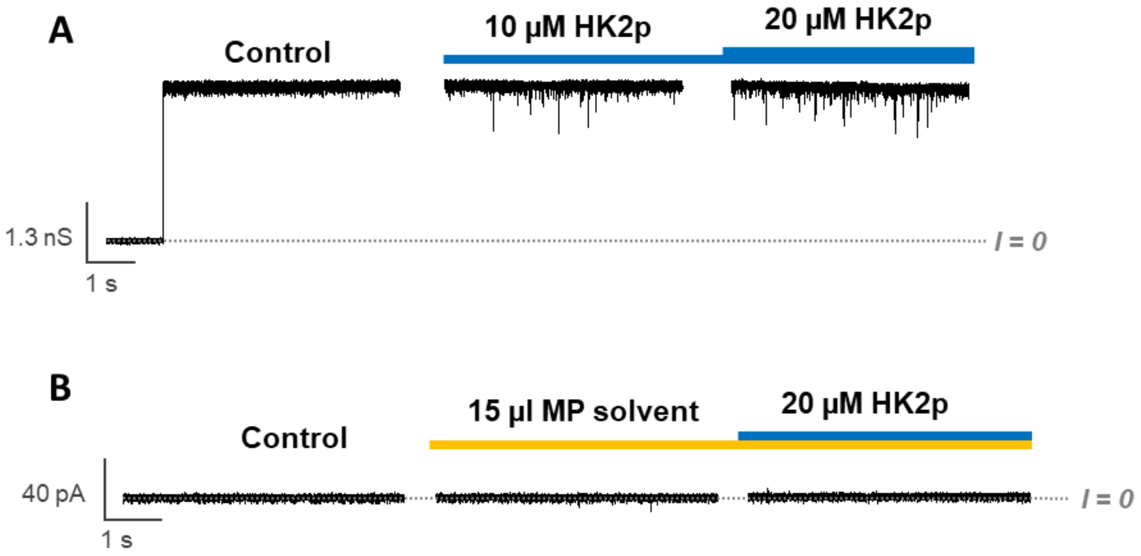


**Supplemental Figure S4.** HK2p induces an excess of current noise through the open VDAC state in PLE/Cholesterol membranes. **(A)** Representative single-channel current record through a VDAC channel before (control) and after the sequential addition of HK2p in MP solvent at 10 µM and 20 µM, as indicated. Consecutive additions of HK2p induce a visible increase of the open-channel current noise. **(B)** Control experiment without reconstituted VDAC, showing that the addition of 15 µl of MP solvent (an equivalent to 20 µM of HK2p addition) or 20 µM HK2p does not cause a leak of the planar membrane or excess of current noise. The planar membrane was formed from PLE/Chol/DPhPC (86/9/5) (mol/mol). The membrane bathing solution contained 1 M KCl solution buffered with 5 mM HEPES at pH 7.4. HK2p and MP solvent were always added to the *cis* side of the membrane. The original current records were digitally resampled-averaged with 10 as a smoothing factor. Applied voltage was -30 mV.

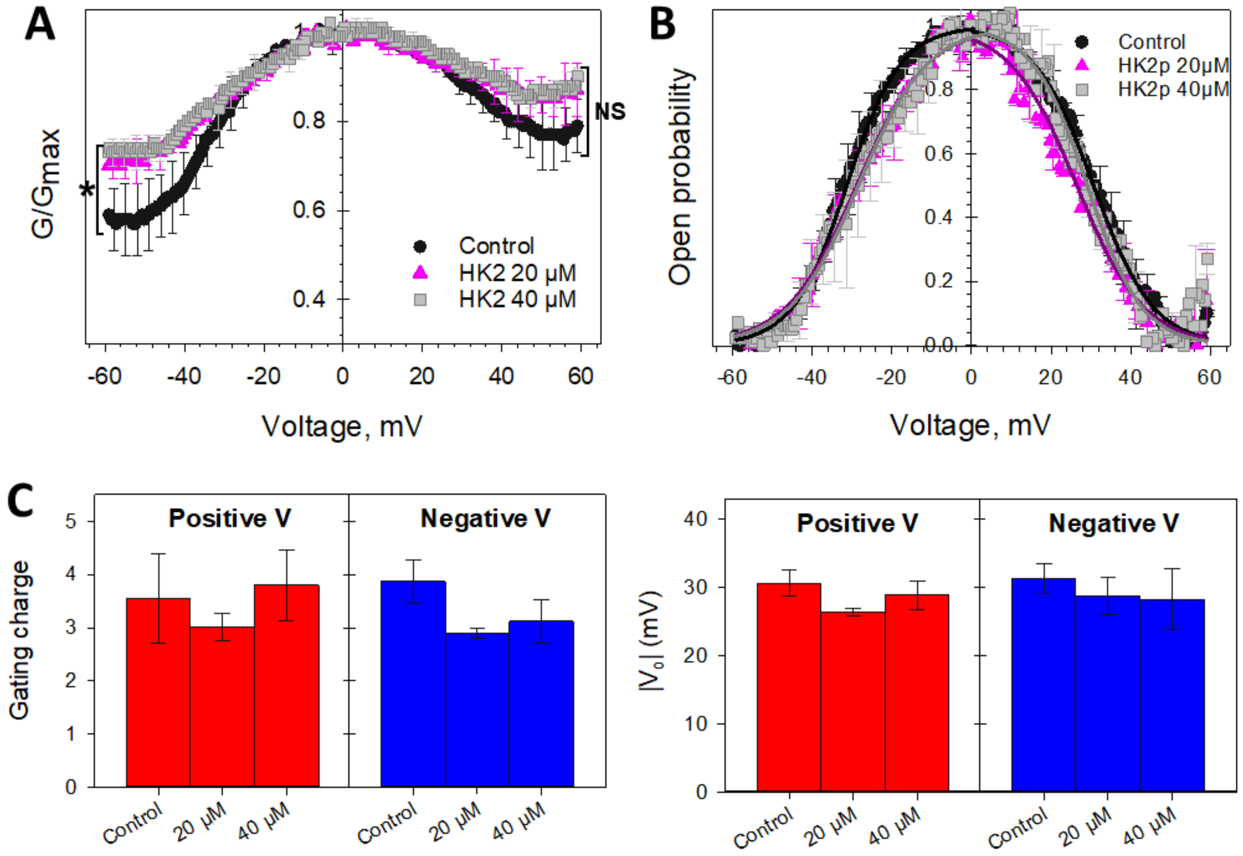


**Supplemental Figure S5.** HK2p moderately decreases VDAC voltage gating. **(A)** Characteristic bell-shaped plot of normalized average VDAC conductance and probability to be open **(B)** as a function of applied voltage before (control) and after consequent addition of 20 and 40 µM of HK2p to both sides of the membrane. Data were obtained on multichannel membranes formed from PLE/cholesterol/DPhPC (86/9/5) (mol/mol) lipid mixture with 10-100 reconstituted VDAC channels. Normalized conductance in (A) is defined as $G/G_{max}$, where $G_{max}$ is the maximum conductance at voltages close to 0 mV. Membrane-bathing solution consisted of 1 M KCl buffered with 5 mM HEPES at pH 7.4. Data are mean of 2-3 experiments and the error bars (shown every 5 points for clarity) indicate the standard deviation from the mean (68 % confidence intervals). Significance (*$p < 0.05$, NS (not significant): $p > 0.05$; Student's t-test) was checked between control and 20 µM HK2p and between 20 and 40 µM HK2p. There is no significant difference between 20 and 40 µM HK2p. The minimum $G/G_{max}$ value of each data set was used, corresponding to $|V| \sim 55$ mV. Each voltage polarity was tested independently. Solid lines in (B) (black for Control, dark pink for 20 µM HK2p, and light grey for 40 µM HK2p) are the fits of $P_{open}$ plots with the Boltzmann equation (Eq. (3)) using the effective gating charge, $n$, and the voltage at which half of the channels are open, $V_0$, as fitting parameters, shown in **(C)**. There is no significant difference between gating parameters $n$ and $V_0$ ($p > 0.05$, Student's t-test). HK2p affects neither the channel open probability nor the gating parameters. Control experiments with additions of the corresponding aliquots of HK2p solvent, MP, showed no measurable effect on VDAC gating.

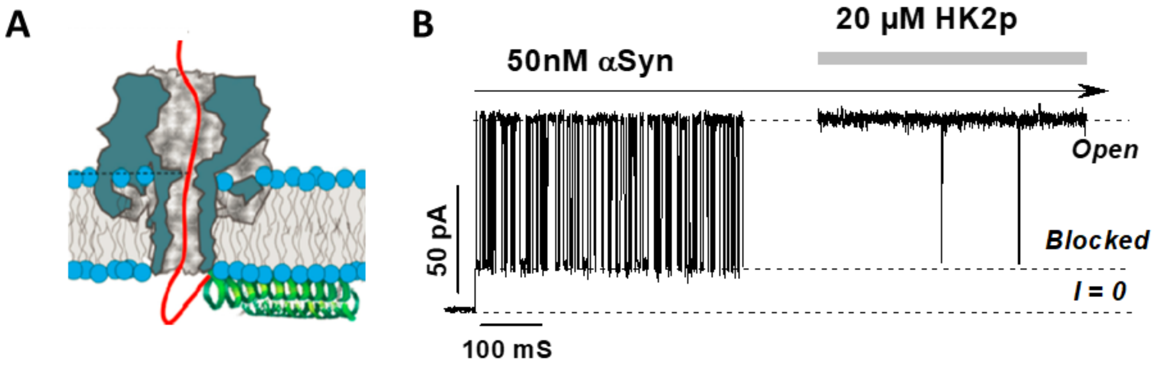


**Supplemental Figure S6.** HK2p inhibits αSyn blockage of α-hemolysin β-barrel channel. (**A**) A cartoon showing α-hemolysin reconstituted into the lipid bilayer and interacting with αSyn. Acidic CTT of αSyn enters the α-hemolysin channel from its stem side (*trans*-side on PLM setup). Adapted with permission from Gurnev et al, *Biophys J.* (2014) [8]. Copyright © 2014 Biophysical Society. (**B**) Representative current record through a single α-hemolysin channel in the presence of 50 nM αSyn in the *trans* side of the membrane and after addition of 20 μM HK2p-TAT to the same side. Single α-hemolysin channel was reconstituted into DPhPC membrane bathed in 1 M KCl buffered with 5 mM HEPES at pH 7.4. Applied voltage was 30 mV. Original current record was digitally filtered at 1 kHz.

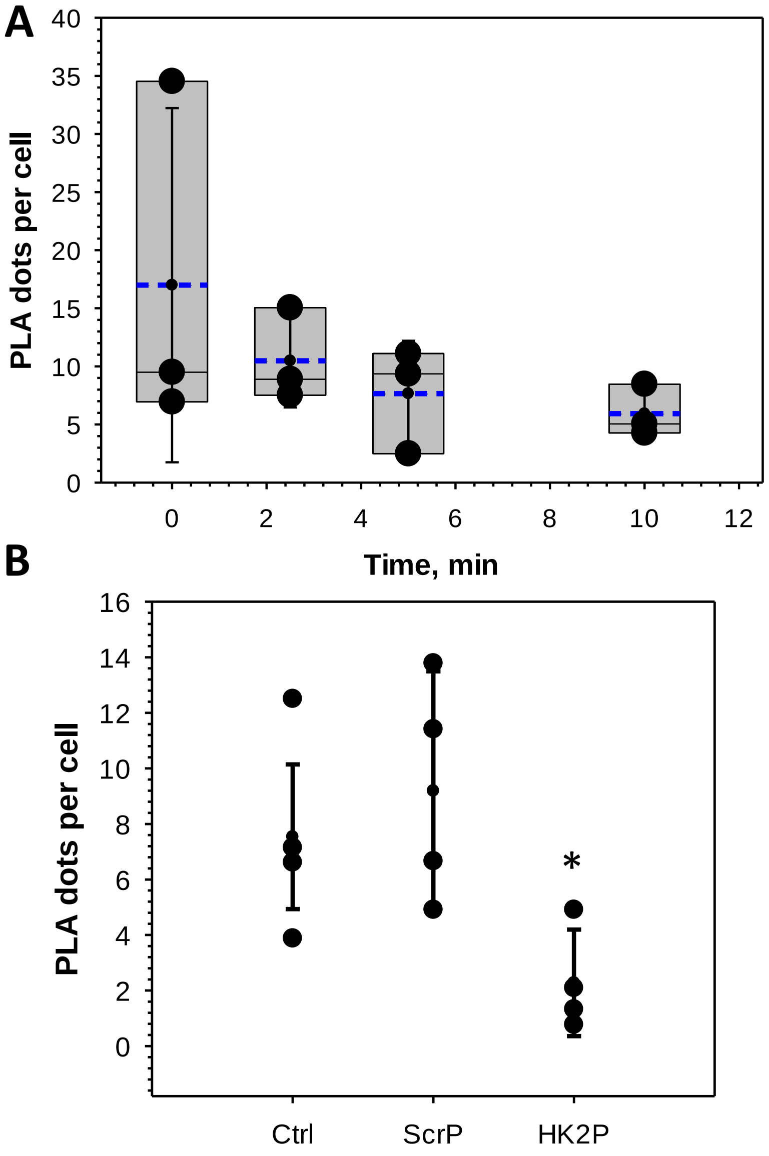


**Supplemental Figure S7.** HK2p reduces αSyn entry into mitochondria. (**A**) Box plots represent the average raw PLA dots per cell at different times after the addition of 5 μM HK2p. (**B**) Box plot of raw PLA dots per cell after treatment with Scrambled (ScrP) or HK2p peptide. The borders of the boxes define the 25$^{th}$ and 75$^{th}$ percentiles, with the dashed blue line as a mean, the median displayed as black lines, and error bars indicate the standard deviation of the mean. The significance was tested against control without treatment using a paired t-test (*$p<0.05$).
